# Molecular characterization of an adhesion GPCR signal transduction

**DOI:** 10.1101/2025.08.14.670383

**Authors:** M. Marfoglia, L. Dumas, B.S. Yang, A. Oggier, M. Pedraza, M. Hijazi, A.N. Larabi, K. Lau, F. Pojer, M.A. Nash, P. Barth

## Abstract

Key cellular processes rely on the transduction of extracellular mechanical signals by specialized membrane receptors, including adhesion G-protein-coupled receptors (aGPCRs). While recent studies support aGPCR activation via shedding of the extracellular GAIN domain, shedding-independent signaling mechanisms have also been observed. However, the molecular basis underlying these distinct activation modes remains poorly understood. Here, we integrate single-molecule force spectroscopy, molecular dynamics simulations, and cell-based assays to elucidate the structural and dynamic mechanisms of ADGRG1 mechanotransduction. We show that shear stress induces distinct deformation pathways in the isolated GAIN domain, promoting tethered agonist (TA) exposure through loop rearrangements prior to domain shedding. In the full-length receptor, defined GAIN orientations and specific loop contacts with the 7-transmembrane (7TM) core enable allosteric TA engagement and signaling in the absence of GAIN dissociation. The directionality of the applied force dictates the activation pathway, favoring either GAIN shedding or intact GAIN–7TM coupling. These mechanisms align with both the basal activity and collagen-enhanced signaling of ADGRG1. Using deep learning-guided design, we engineered GAIN variants with tailored mechanical sensitivity, validating our model through predictable shifts in constitutive and ligand-induced signaling. Together, our findings establish a unified framework for aGPCR activation governed by GAIN dynamics and orientation, bridging mechanical and allosteric models of receptor function and providing new strategies for engineering mechanosensitive receptors and precision therapeutics.

## Introduction

Cellular mechanosensing is currently thought to primarily involve integrins, which couple the extracellular matrix (ECM) to the contractile cytoskeleton^1–3^, or mechano-activated ion channels^4, 5^. However, adhesion GPCRs (aGPCRs) bearing large extracellular regions (ECR) capable of binding to ECM components^6–8^ have received growing attention due to their role in mechanical regulation^9–13^. Virtually all receptors in this family bear a highly conserved GPCR-Autoproteolysis INducing (GAIN) domain located in the ECR. Typically situated between the adhesion ligand binding domain (LBD) and the signaling 7 transmembrane (7TM) domain, the GAIN structure carries a tethered peptide agonist (TA or Stachel) that often undergoes auto-proteolytic cleavage but remains deeply buried inside the GAIN structure^6, 14^. aGPCR signaling under mechanical stress can occur upon GAIN dissociation from the TA^15^. With GAIN dislodged, the TA then binds the 7TM domain and triggers intracellular signaling^7, 16, 17^. Recent cryoEM structures of shedded aGPCRs in active signaling states (i.e. bound to GAIN-dissociated TA and G-proteins) and single-molecule force spectroscopy on isolated GAIN domains support this model of receptor activation through GAIN shedding^18–22^. However, several lines of evidence support an alternative allosteric mechanism of receptor activation that does not require GAIN dissociation^6, 16, 17, 23–25^. In absence of high-resolution full length aGPCR structures, the molecular basis of these distinct allosteric and shedding mechanisms of activation remains unclear.

Protein structures can experience significant deformations upon mechanical loading, resulting in force-activated phenotypes and resistance to applied stress^26, 27^. Molecular deformation can alter the orientation of covalent and non-covalent bonds within the protein structure, and represents a key determinant of protein mechanical response^26–28^. In recent years, proteins with novel shapes and topologies have been designed *de novo* using deep-learning approaches trained on static structures^29^. Engineering powerful mechano-sensing functions would require understanding protein structures and dynamic adaptations under mechanical stress, a concept that has not yet been exploited in protein design.

In this study, we aimed to uncover how GAIN interacts with the 7TM domain and mediates signal transduction under distinct chemical and mechanical stimuli (**Fig. 1A**). We tackled this question using a combination of single-molecule force spectroscopy, cell signaling assays, molecular simulations and a novel deep learning approach that investigates protein motions under mechanical load^30^. Our multidisciplinary approach revealed novel structural and dynamic principles of aGPCR activation at unprecedented resolution.

**Figure 1.**
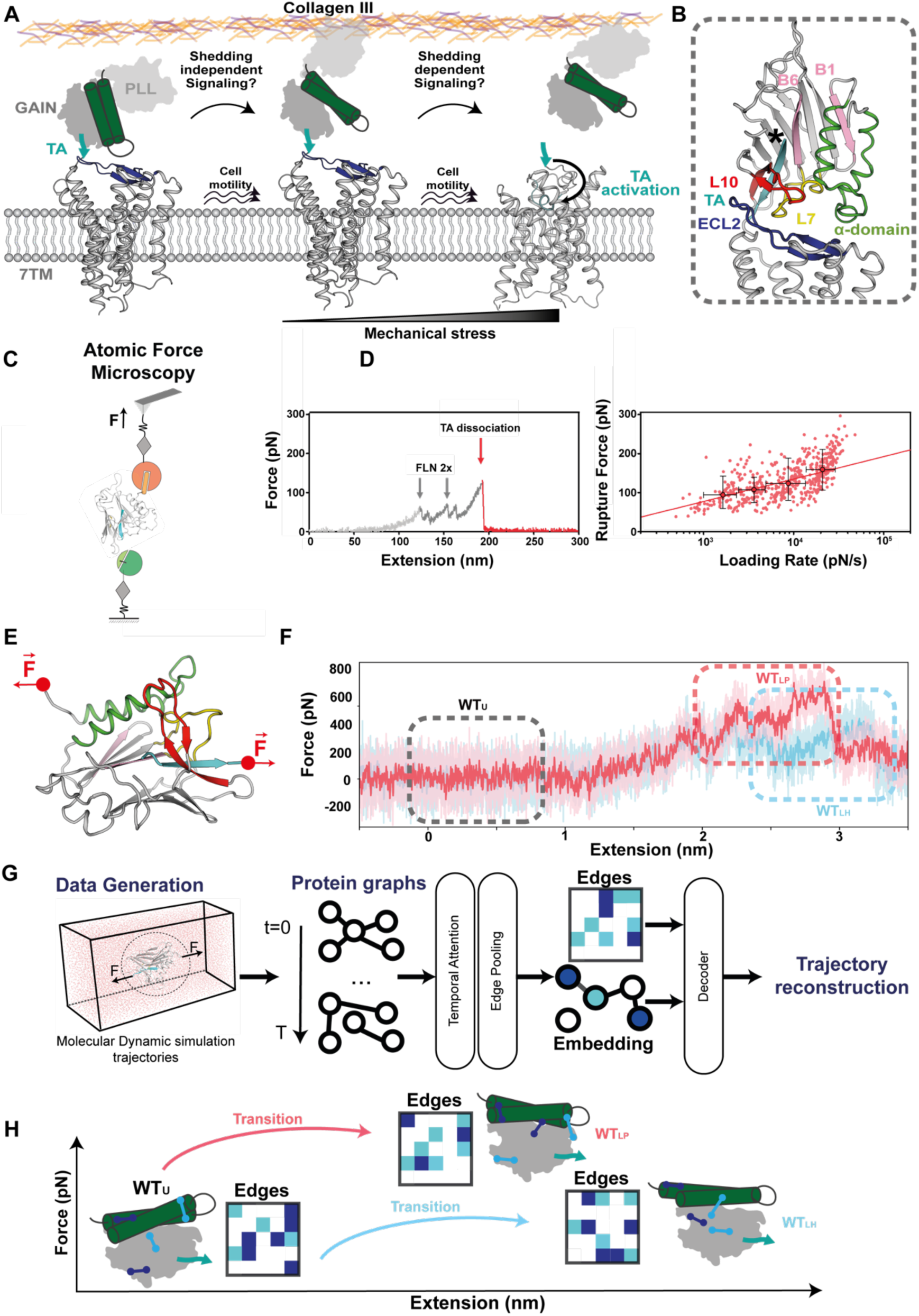
The Adhesion GPCR GAIN-TA interface behaves as a stable mechanical clamp motif. **A.** Putative activation mechanisms of ADGRG1 as a function of mechanical shear stress, where ADGRG1 signaling can be mediated through shedding-independent or shedding-dependent mechanisms. **B.** Structural model of ADGRG1 obtained by Alpha-Fold showing extensive contacts between flexible regions of the GAIN domain (Loop 1: green, Loop 7: yellow, Loop 10/Lid: red, Tethered Agonist (TA): cyan) and the 7TM extracellular region (ECL2: blue). **C,D.** Force-extension curve and dynamic force spectrum of the GAIN domain characterized by AFM-SMFS. **D.** Rupture forces fall in a range of 100-150 pN at the indicated loading rates (∼10^4^ pN/s). **E,F.** Steered Molecular Dynamic (SMD) simulations of the GAIN domain performed by pulling from its N- and C-termini (**E**) with representative force-extension curves (**F**) denoting the variability in mechanical response observed in the trajectories and defining two distinct loaded states: polar loaded (WT_LP_) and hydrophobic loaded (WT_LH_) states, respectively. **G.** Architecture of the AlloPool Graph Neural Network (GNN) model^30^, designed to learn the minimal set of tertiary contacts (edges) required to predict protein conformational dynamics from molecular dynamics simulation trajectories. Timeseries of protein states extracted from MD simulation trajectories serve as input to an auto-encoder which learns key edges regulating the protein dynamics. **H.** Schematic representation of the distinct sets of tertiary contacts (edges) that govern different conformational transitions of GAIN during SMD. Interactions between residue pairs are depicted as boxes on a 2D edge map, where each axis represents the protein sequence. Edges are color-coded by strength (weights): dark blue (high), light blue (medium), and white (low).

## Results

We selected the canonical and well-studied aGPCR ADGRG1 receptor because it bears a topologically simple ECR composed of one N-terminal ligand-binding domain (PTX/LNS-Like, PLL) linked to the 7TM signaling region through an autoproteolytically cleaved GAIN domain (**Fig. 1A**). The structure of the isolated murine homolog ADGRG1 EC region was solved by X-ray crystallography^31^. The cryoEM structure of the 7TM region in an active signaling conformation bound to the GAIN-dissociated TA and G-protein was released recently^18^. In the absence of an experimentally-characterized full length ADGRG1 structure, we built a model using AlphaFold from the structures of each domain to obtain a coarse-grained view of the GAIN-7TM interface. The model revealed 3 main regions of the GAIN domain potentially mediating important interactions with the 7TM domain. These include loops L1, L7 and L10 (the so called Lid) that either connect the alpha-domain or stabilize the beta-sheet region where the TA is buried (**Fig. 1B, Extended Data Fig. 1**).

### ADGRG1 GAIN displays high mechanical resistance

Before investigating how GAIN mediates mechanotransduction through these flexible regions in the context of the full-length ADGRG1 receptor, we first sought to understand how the GAIN structure responds to mechanical forces at the single-molecule level. Previously, a single-molecule magnetic tweezers (MT) study investigated the mechanical properties of GAIN and demonstrated the possibility for force-mediated shedding, but did not reveal significant mechanistic and structural insights into GAIN responses to mechanical stress^32, 33^. Here we characterized the mechanical properties of the ADGRG1 GAIN domain using atomic force spectroscopy (AFM) to provide complementary analysis to MT and enable comparison to other mechanical proteins (e.g. integrins, titins etc…) studied by AFM^33–36^. It is known that a protein’s mechanical response is highly dependent on the loading point and direction of the applied force^28, 37–42^. In the context of the full-length ADGRG1 receptor expressed at the cell surface, mechanical force is likely transduced to GAIN by the N-terminal adhesion PLL domain bound to an ECM ligand. To best mimic this native orientation and direction of the applied force in our AFM experiments, we designed a setup where the C-terminal end of GAIN was covalently attached to a solid surface while the pulling force was applied to the N-terminal end of the domain through a regeneratable peptide/receptor complex (FgB/SdrG) (**Fig. 1B, Extended Data Fig. 2, Methods**). This experimental design enabled fresh GAIN molecules to be probed on the surface and allowed repeated measurements and quantification of TA dissociation events by AFM. We measured WT GAIN-TA dissociation events in the range of 100-160 pN with a clear dependence on the loading rate over a range from 1,000 – 40,000 pN/s (**Fig. 1B-C**). These results indicate that the GAIN-TA interface displays relatively high mechanical stability on par with integrins but substantially higher than those measured for the Notch receptor^43–47^. When we extrapolated our data to equivalent loading rates used in the MT study (1 pN/sec), we obtained similar rupture forces in the range of 20 pN, suggesting that the discrepancies between MT and AFM-derived values can largely be accounted for by different experimental measurement conditions.

### GAIN behaves as a multiswitchable mechanosensor

To investigate how the GAIN structure responds to mechanical forces and elucidate which structural features might confer mechanostability to the GAIN-TA interface, we performed Steered Molecular Dynamics (SMD) simulations by applying a constant harmonic force potential to the C-terminal structure and pulling simultaneously from the N terminus of the GAIN to mimic the force directionality in AFM (**Fig. 1D**). We ran simulations at pulling speeds as low as 0.1 nm.ns^−1^. While this speed is orders of magnitude faster than those used in AFM, SMD performed at such rates were shown to recapitulate general unfolding and unbinding events measured by AFM^26^. Pulling on the protein from its N- and C-termini resulted in an increasing amount of force accumulated within the structure, reaching high mechanically resistant (i.e. loaded) states prior to unfolding and TA dissociation (**Fig. 1E**). Interestingly, the SMD trajectories were not uniform and revealed a complex behavior where the GAIN structure accessed multiple loaded states with distinct levels of mechanical resistance. Extracting physiologically-relevant structural transitions and motions from heterogenous atomistic MD simulations remains very challenging as the protein explores multiple time-dependent states and structural fluctuations are often dominated by random noise. To address that problem, we developed AlloPool, a dynamic Graph Neural Network-based approach for inferring protein conformational dynamic properties from heterogenous MD simulation datasets^30^ (**Fig. 1F**, **Methods**). The method learns from multiple simulation trajectories the set of atomic interactions (i.e. pooled edges in protein graphs, **Fig. 1F,G**) that govern distinct time-dependent protein structural transitions and allosteric motions (**Fig. 1G, Methods**). Unlike alternative approaches^48^, AlloPool can predict complex molecular motions over long timescales^30^ (**Methods**).

When applied to the SMD trajectories of the GAIN, AlloPool identified transitions to two main loaded states (WT_LP_ and WT_LH_), whose distributions depended on the pulling rate of the SMD simulations (**Fig. 2A**). These transitions define distinct GAIN responses to mechanical stress and involve the breaking and formation of different sets of tertiary structural and dynamic contacts between the alpha sub-domain (A), loops and GAIN-TA interface (**Fig. 2B-G**). In the hydrophobic loaded (i.e. WT_HL_) state mainly observed at high pulling speeds, the alpha-domain aligns colinearly to the force. Mechanical resistance is mostly achieved through enhanced hydrophobic packing contacts in the protein core mediated by the closure of L7 and L10 onto the alpha sub-domain (**Fig. 2B,D,F**). In this compact state, the hydrophobic core sustains the force and the TA remains largely buried in the GAIN scaffold.

**Figure 2.**
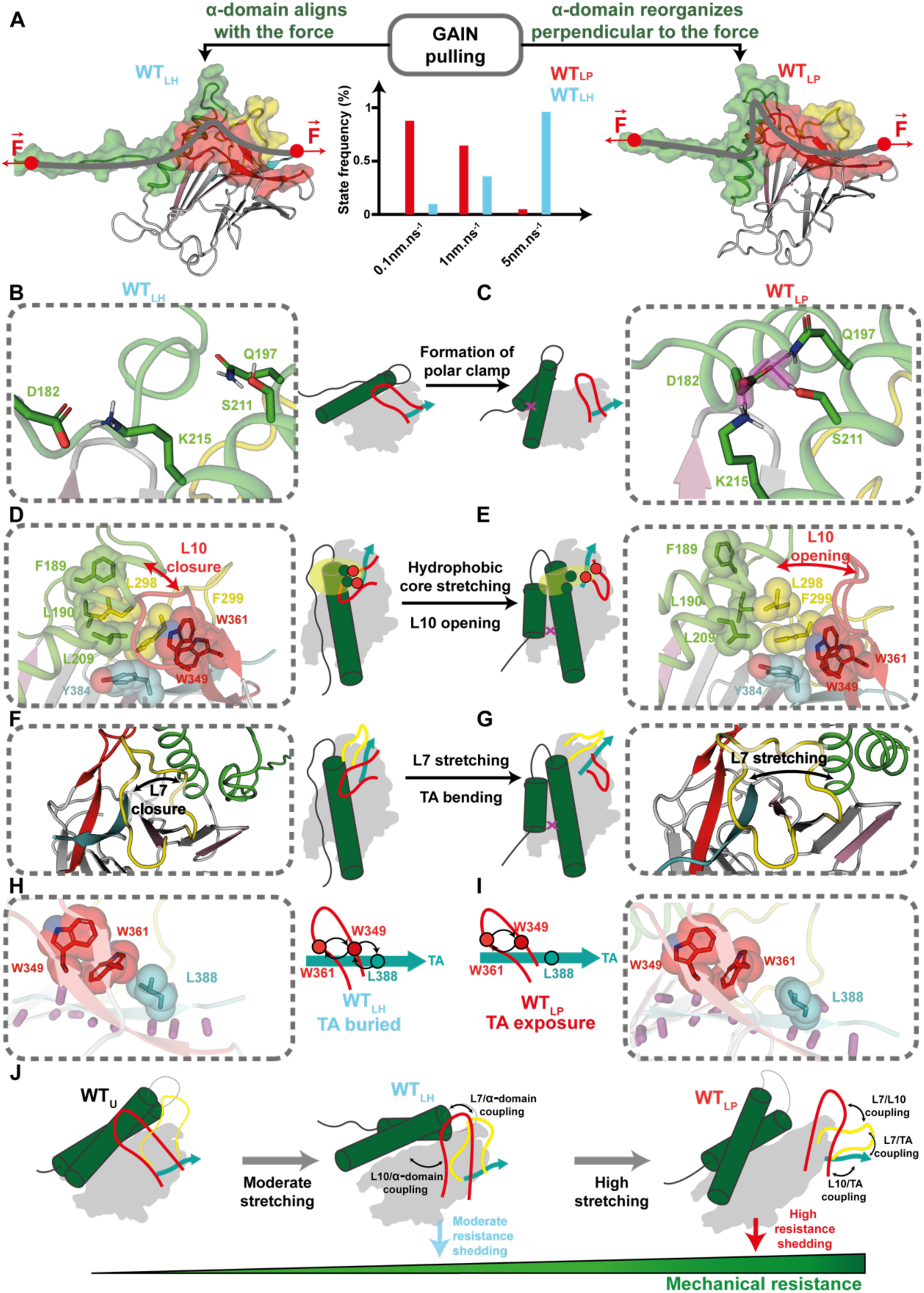
Structural reorganization of the GAIN domain upon mechanical loading. **A.** GAIN accesses two structurally distinct mechanically loaded states: hydrophobic loaded: WT_LH_ (left) and polar loaded: WT_LP_ (right) whose distribution depends on the pulling speed of the simulations. The α-domain aligns either colinearly or perpendicularly to the pulling force in the WT_LH_ and WT_LP_ states, respectively. GAIN reaches higher mechanical resistance in the WT_LP_ state. **B,C.** The α-domain undergoes large structural reorganization in the WT_LP_ state enabling the formation of a strong non-native polar network stabilizing the interface between helices 1 and 2 (i.e. polar clamp) (**C**). This clamp is not observed in the WT_LH_ state. **D,E,F,G.** While the WT_LH_ state achieves mechanical resistance through enhanced hydrophobic packing between helix-1, L7 and L10 that closed on the α-domain (**D,F**), GAIN in the WT_LP_ state undergoes hydrophobic core stretching (**E,G**) and L10 opening that tightly coupled with the TA. **H,I.** Structural reorganization of the GAIN domain in the WT_LP_ state triggered bending, partial dislodging from the hydrophobic core and enhanced solvent exposure of the TA (**I**). TA remained buried within GAIN in the WT_LH_ state. **J.** The GAIN domain senses and responds to mechanical shear stress through a gradient of structural deformations to a wide range of applied mechanical pulling conditions. Structural reorganization of GAIN involves conformational changes and differential coupling between switchable motifs (loops L7, L10, TA and α-domain). GAIN achieves different levels of mechanical resistance through distinct motions and coupling of these switchable motifs.

The alternative polar loaded (i.e. WT_PL_) state occurs at lower pulling speed, involves larger conformational changes and reaches higher mechanical resistance than WT_HL_. In WT_PL_, the alpha domain moves away from the rest of the scaffold, perpendicular to the applied force, and undergoes interhelical reorientation allowing the formation of a strong polar catch bond between the helices (**Fig. 2A,C**). The alpha domain motions stretch the hydrophobic core and trigger the opening of L10 and L7 which partially dissociate from the alpha domain. The 2 loops can then fold back onto the TA and enhance the coupling of the GAIN-TA interface to withstand higher mechanical shear stress (**Fig. 2E,G**). This reorganization ultimately leads to the bending of the TA strand which becomes partially exposed to the solvent (**Fig. 2I**).

Overall, GAIN acts as a multi-state switchable mechanosensing domain, where coordinated motions and interactions between local switchable regions (H1-H2, L7, L10, and TA) create a gradient of allosteric responses to different levels of applied mechanical stress (**Fig. 2J**). At high pulling rates, GAIN undergoes limited stretching along the N-C terminal axis, primarily reinforcing the hydrophobic packing that provides moderate mechanical resistance. At lower pulling rates, GAIN can undergo greater stretching and structural adaptations, including the formation of polar clamp motif. While these motions further enhance mechanical resistance, they also lead to partial dislodging and solvent exposure of the TA, potentially impacting GAIN-7TM interaction and signal transduction.

### Designed GAIN domains with reprogrammed motions and mechanical stability

Overall, AlloPool uncovered key structural determinants governing GAIN responses to mechanical stress including several regions predicted to interact with the signaling 7TM domain in structural models of the receptor (**Fig. 1A**). To stringently test these predictions, we leveraged AlloPool’s predictions to engineer GAIN variants with reprogrammed structural responses to mechanical stress. We developed a deep-learning-based computational framework to design proteins with customized mechanical stability (**Fig. 3A**). This method integrates structure-based design with AlloPool-guided motion prediction to engineer novel sequence-structure motifs and conformational dynamic behavior under a broad range of mechanical stress conditions.

**Figure 3.**
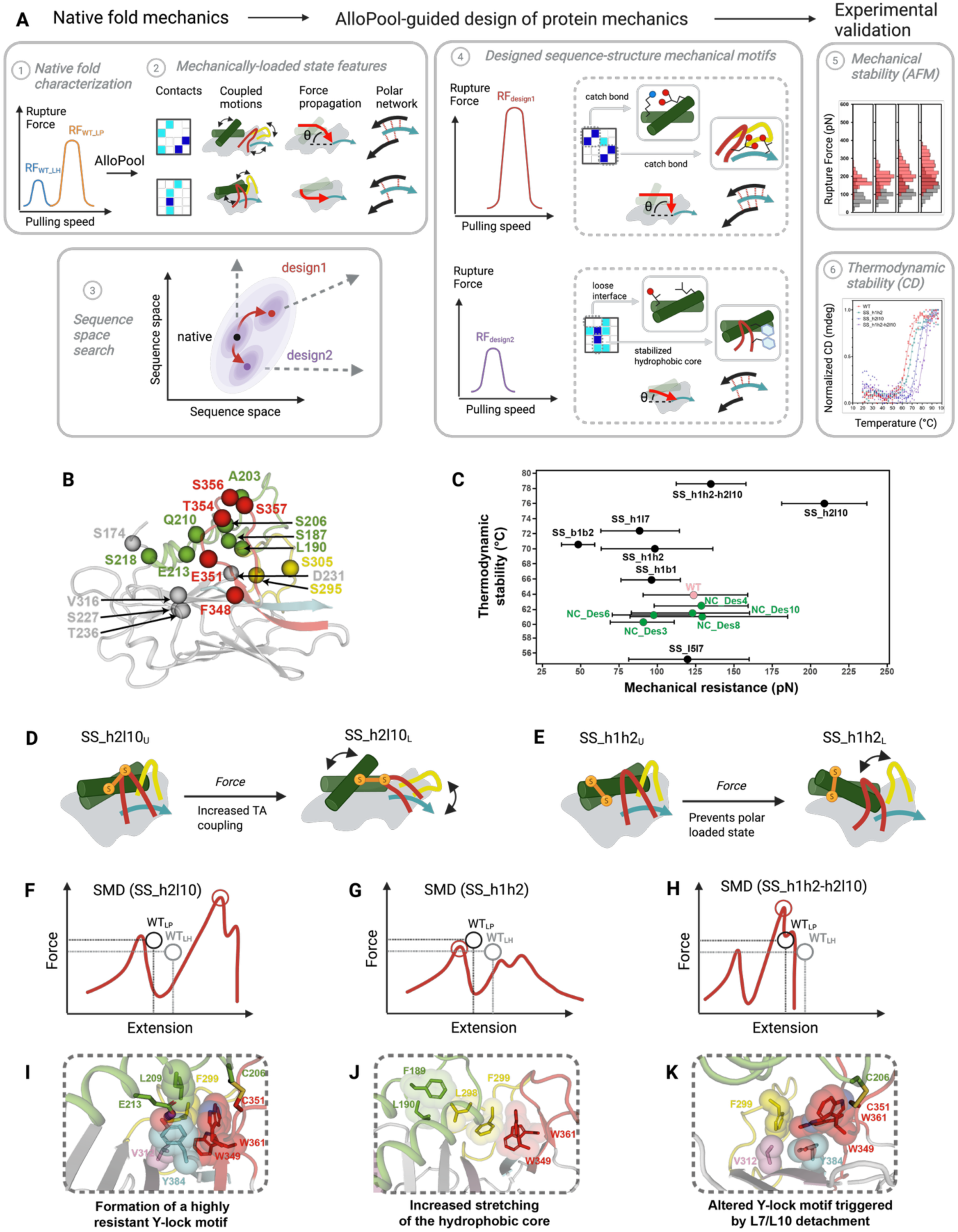
Computational design of GAIN domains with reprogrammed motions and mechanical resistance. **A.** Computational flowchart for the Deep Learning (DL)-guided design of protein mechanical properties (Methods): 1,2. Characterization of the sequence-structure underpinnings of a native fold mechanical stability. 3. Search for designed sequences reprogramming fold mechanical stability. 4. In silico characterization of designed sequence-structure mechanical motifs. 5. Experimental validation of thermodynamic and mechanical stability. **B.** Positions of the designed residues mapped on the 3D structure of GAIN. **C.** Experimentally-measured mechanical and thermodynamic stabilities of the designed GAINs. SS and NC denote domains with designed disulfide bridges or non-covalent interaction networks, respectively. Horizontal bars represent the range of rupture forces measured at distinct pulling speeds. WT GAIN is represented in pink. **D, E.** Design goals of disulfide locks engineered using interaction patterns learned from AlloPool. Expected impacts of the disulfide bridges on the unloaded (U) to loaded (L) state transition of the GAIN SS_h2l10 (**D**) and SS_h1h2 (**E**) variants. **F-H.** Schematic description of the SMD force-extension curves (red dotted lines) obtained for the SS_h2l10 (**F**), SS_h1h2 (**G**) and SS_h1h2-h2l10 (H) designed variants as compared to the WT_LP_ (dark gray dotted line) and WT_LH_ (light gray dotted line) curves. Loaded states are shown by a circle with associated key motifs identified by AlloPool and responsible for the observed mechanical resistance. **I-K.** Structural view of the mechanically-resistant motifs in the loaded states for SS_h2l10 (**I**), SS_h1h2 (**J**) and SS_h1h2-h2l10 (**K**).

The framework operates through a multi-step process. First, it characterizes the mechanical phenotype of a given protein fold by identifying key determinants of mechanical stability. These include the specific network of contacts and conformational changes that govern the transitions to mechanically loaded states, the interactions stabilizing the rupture interface—such as hydrogen bond networks at the GAIN-TA interface—and the force propagation pathways that determine how effectively an applied force disrupts a structural interface^26, 28^ (**Extended Data Fig. 3**). Next, the method conducts a multi-state search to identify sequence-structure motifs and dynamic properties that selectively modulate these mechanical features while preserving optimal thermodynamic stability in the resting state (**Methods**). This enables the engineering of novel catch bonds and tailored conformational changes to enhance mechanical resistance. Alternatively, it can be used to weaken particular protein interfaces or redirect force transmission, thereby reducing mechanical stability. Finally, designed protein variants exhibiting the intended mechanical behavior undergo experimental validation through biophysical measurements. Thermodynamic stability is assessed using techniques such as Circular Dichroism in bulk solution, while mechanical stability is evaluated under mechanical tension using Atomic Force Microscopy (AFM).

We applied this approach to the GAIN domain and identified several variants with modified tertiary contact networks and disulfide-bridge-mediated covalent bonds in the hydrophobic core, α/β-sheet interfaces, and switchable loops L5, L7 and L10. These modifications reprogrammed GAIN coupling, motions and force propagation, giving rise to novel switchable behaviors and mechanically resistant loaded states (**Fig. 3B-K**).

A representative subset of 12 promising variants was selected for experimental characterization. This included designs featuring novel non-covalent interaction networks spanning the α-subdomain, hydrophobic core, and loop L10 (e.g., NC_Des3, NC_Des4, NC_Des6, NC_Des8 and NC_Des10). Additionally, we identified variants incorporating designed disulfide bridges predicted to stabilize switchable regions, including helices h1 and h2, β-strands 1 and 2, and loops L5, L7 and L10. Our design objective was to modulate mechanical response while maintaining thermodynamic stability in non-covalent designs, whereas properly formed disulfide bridges were expected to enhance the thermodynamic stability of the folded ground state. To evaluate this, we measured the thermal unfolding of the GAIN domain via circular dichroism and found that all non-covalent designs retained thermodynamic stability comparable to wild-type (Tm > 60°C). Except for SS_l5l7, all disulfide-bridge-based designs exhibited enhanced thermodynamic stability, consistent with correct covalent bond formation (**Fig. 3C, Extended Data Fig. 4**). Next, we assessed the mechanical properties of these variants. Using identical functionalization and experimental conditions as WT, we recorded hundreds of atomic force microscopy (AFM) force-extension curves, filtering for single-molecule interactions and quantifying GAIN-TA dissociation forces. The resulting rupture forces spanned a broad range, from as low as 67–117 pN to as high as 182–236 pN (**Fig. 3C, Extended Data Fig. 5**). These data aligned well with our design goals and confirmed that thermodynamic and mechanical stability are distinct properties of the protein’s ground and excited states, which can be engineered independently.

An in-depth analysis of the designed features demonstrated excellent agreement between AlloPool’s mechanistic predictions and the measured mechanical phenotypes. For this analysis, we selected two non-covalent designs (NC_Des3, NC_Des6) that exhibited weaker mechanical resistance than WT, along with three topology-rewired variants: SS_h1h2, SS_h2l10, and SS_h1h2_h2l10. These variants displayed the lowest (67-135 pN), highest (182-236 pN), and no significant difference (115-158 pN) in mechanical resistance compared to WT, respectively (**Fig. 3C**). Additionally, we solved the structure of the double cross-linked SS_h1h2_h2l10 variant using X-ray crystallography, which closely aligned with our structural models (Cα RMSD = 2 Å) and confirmed the conformation of the engineered disulfide bridges (**Methods**, **Supplementary Discussion**, **Extended Data Fig. 6-8, Extended Data Table 1**).

By locking L10 to helix 2, SS_h2l10 enhanced the mechanical resistance of the GAIN-TA interface by increasing coupling between TA and L7, as observed in the WT_PL_ loaded state (**Fig. 3D**, **Extended Data Fig. 9-10**), while maintaining tight connections between TA and L10. Unlike in WT, where only one β-strand and L7 are dynamically coupled to TA, all components of the TA-GAIN interface in the SS_h2l10 variant are strongly coupled. This ensures an optimal distribution of mechanical force across the entire interface, thereby enhancing its mechanical resistance— similar to what was observed in the SdrG-FgB interface^26^. In our SMD simulations, we observed the formation of an initial mechanically resistant structure resembling WT_LP_, followed by a second state exhibiting a 2.5-fold increase in force resistance compared to WT (**Fig. 3F, Extended Data Fig. 11**). This newly identified loaded state is characterized by a highly stable GAIN-TA interface, where the highly conserved Y384 of the TA is locked through buried polar and hydrophobic interactions with a cluster of residues on helix 2, L7, and L10 (i.e., the Y-lock motif, **Fig. 3I**). Additionally, this state exhibits several features reminiscent of highly mechanically stable interfaces (**Supplementary Discussion**, **Extended Data Fig. 10, Extended Data Fig. 12)**.

Mechanically weaker GAIN variants were generated by either locking helices 1 and 2 through a disulfide bridge (SS_h1h2 variant, **Fig. 3E**) or reinforcing the hydrophobic core via multiple non-covalent interaction networks (NC_Des3, NC_Des6 variants). These designs prevented the formation of the WT_PL_ polar clamp motif between helices 1 and 2, forcing both helices to align collinearly with the applied force, rather than only helix 1 as observed in WT. Additionally, they maximized the force exerted on the hydrophobic core and encoded several features predictive of mechanically looser interfaces (**Supplementary Discussion, Extended Data Fig. 10, Extended Data Fig. 12**). In the topologically constrained SS_h1h2 scaffold, the loaded state is reached more rapidly but exhibits mechanical resistance similar to WT_HL_ (**Extended Data Fig. 11**). During this state, the hydrophobic core undergoes significant stretching and is disrupted shortly after the force peak (**Fig. 3G, J**).

Lastly, we characterized the double mutant SS_h1h2_h2l10, which combines both designed disulfide bridges. In our simulations, this variant initially behaves similarly to SS_h1h2 before transitioning to a more mechanically resistant state, stabilized by a partial Y-lock motif (**Fig. 3K**). However, the presence of both disulfide locks induces extreme stretching of the alpha-domain, leading to the decoupling of L7 and L10. This disruption prevents the formation of an optimal Y-lock interaction network and weakens TA coupling with neighboring β-strands, ultimately making it less mechanically resistant than SS_h2l10 (**Fig. 3H,K, Extended Data Fig. 10-12**, **Supplementary Discussion**).

The strong agreement between AlloPool’s predictions and the measured mechanical responses of the designed GAINs validates our molecular mechanistic description of GAIN structural responses to mechanical forces. Moreover, it highlights that protein mechanics can be engineered by reprogramming the long-range dynamic properties of the protein scaffold, as identified under mechanical load using our novel deep-learning approach, AlloPool.

### ADGRG1 signals through allosteric coupling between the 7TM and the GAIN in absence of mechanical shear stress

With distinct TA rupture forces and dynamic properties in regions contacting the signaling 7TM domain, both native and designed GAIN domains provide a unique opportunity to deepen our understanding of the molecular mechanisms underlying ADGRG1 activation. Structural characterization of full-length adhesion GPCRs has been highly challenging, largely due to the flexible nature of their extracellular regions. To date, only very low-resolution maps of the GAIN-7TM interface have been obtained^25^, hindering our comprehension of aGPCR signal transduction at the molecular level.

To address this challenge, we employed a hybrid computational-experimental approach for characterizing the dynamic properties of the GAIN-7TM interface at atomic resolution and evaluating the behavior of switchable GAIN regions within the context of the full-length receptor. Initially, we conducted 2-microsecond-long equilibrium MD simulations of full-length ADGRG1 receptors using structural models built with AlphaFold based on our X-ray structure of GAIN and the cryo-EM structure of the 7TM domain.

Although the initial GAIN and 7TM structures are in their resting and active signaling states, respectively, the relatively long simulation timescale allows both domains to explore a wide structural space and adopt various conformations that reflect their mutual interactions (**Methods**, **Extended Data Fig. 13**). AlloPool analysis of WT receptor simulations identified three distinct conformational ensembles: a highly flexible state where GAIN remains largely dissociated from the 7TM region, and two compact states—TA-uncoupled and TA-coupled—where GAIN strongly interacts with the 7TM through distinct binding interfaces (**Fig. 4A, B**). Notably, the receptor’s conformational space does not form a continuum but rather consists of well-defined discrete states, closely mirroring recent low-resolution cryo-EM and smFRET findings on ADGRL3^25^.

**Figure 4.**
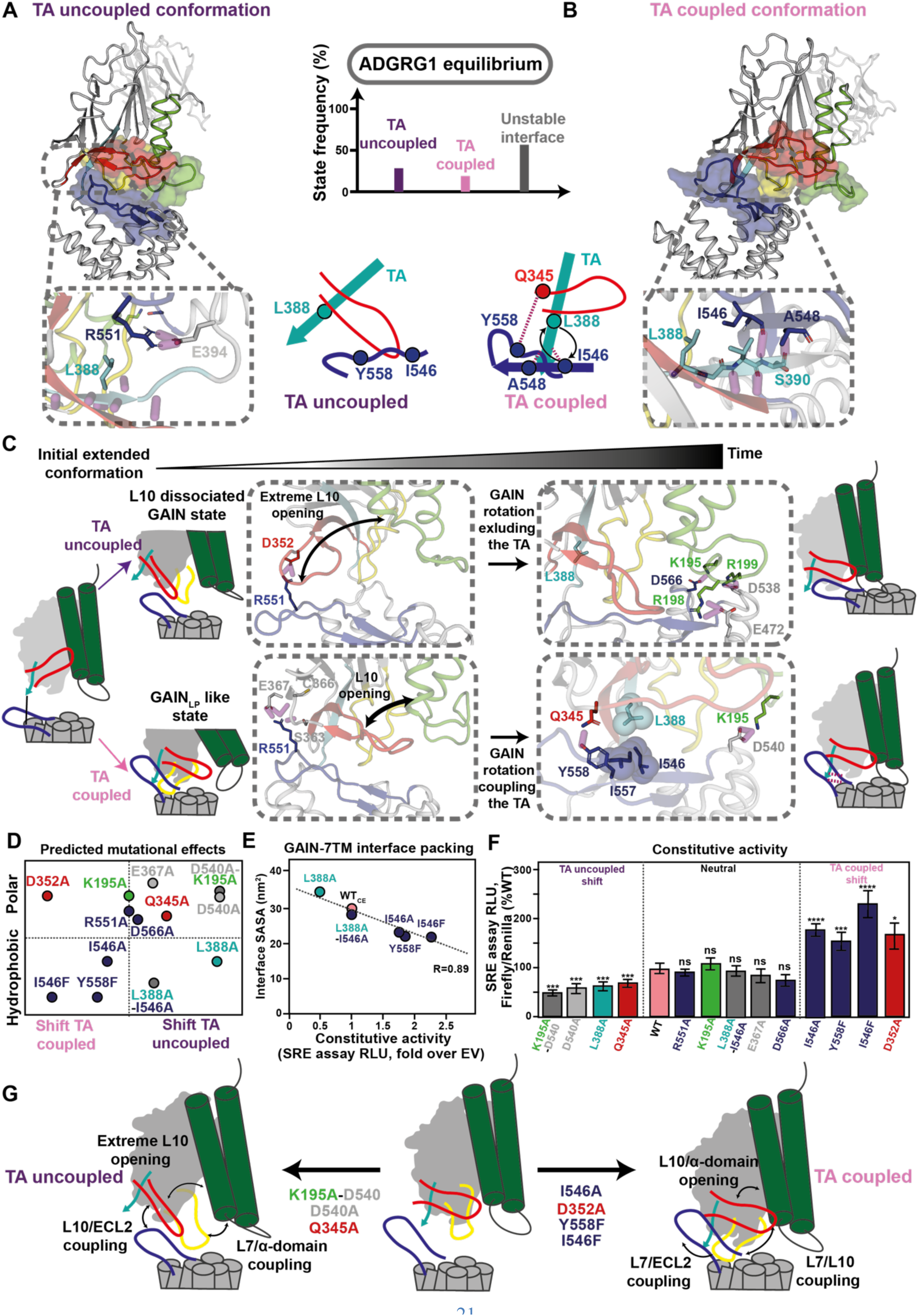
ADGRG1 basal signaling activity is achieved through allosteric coupling between the GAIN and the 7TM domains. **A-B.** Representative structures and schematic representations of TA-uncoupled (**A**) and TA-coupled (**B**) state, with associated frequencies observed in equilibrium MD simulations. In the TA-uncoupled state (**A**, zoom), the TA is excluded from the interface and does not contact ECL2. In the TA-coupled state (**B**), TA directly interacts with ECL2 through 2 backbone hydrogen bonds with I546 and A548. **C.** Conformational changes occurring during the transition between the extended state and the TA uncoupled (top) and TA coupled states (bottom). Middle panels: intermediate conformations towards the TA-uncoupled and TA-coupled states, showing the difference in L10 dynamics, that considerably opens in the TA-uncoupled state (top), as opposed to a WT_LP_-like opening for the TA-coupled state (bottom). **D.** 2D map of ADGRG1 mutants plotted according to the predicted shift towards TA-coupled or uncoupled state and the type of altered interactions (polar or hydrophobic). Mutations are color-coded according to their respective regions within the GAIN domain. **E.** Changes in GAIN-7TM interface packing measured by the interface solvent accessible surface area (SASA) differences correlates with the measured constitutive activity (R=0.89). **F.** Constitutive activity of the designs measured by SRE-luciferase assays. Designs are arranged according to their predicted effects described in **D**. Significant differences to WT activity were calculated by one-way ANOVA and are denoted by stars (***: p-value < 0.001; ns: non-significant). All variants displayed similar cell surface expression (**Extended Data Fig. 15**). **G.** Schematics of the motions following the formation of the TA-uncoupled (left) and TA-coupled (right) states with associated mutations shifting ADGRG1 equilibrium towards specific signaling state.

In these two compact states, the orientation of GAIN and specifically L7, L10, and TA relative to the extracellular loop 2 (ECL2) of the 7TM domain is different, impacting signal transduction. Specifically, in the TA-uncoupled state, the TA is excluded from the ECR-7TM interface and does not contact the extracellular loops (**Fig. 4A**), while in the TA-coupled state, it is well-positioned to interact with ECL2 through hydrophobic contacts and hydrogen bonds bridging the TA and ECL2 beta-strands (**Fig. 4B**). AlloPool revealed that allosteric communication between GAIN and the 7TM is stronger when the TA is partially exposed and engaged with ECL2 (**Extended Data Fig. 14**). Hence, the TA-coupled state likely represents the most signaling-competent conformation of ADGRG1 in the absence of ligand and mechanical stimulus, providing key potential atomic resolution insights into the receptor’s significant constitutive activity (see below).

To validate these findings, we identified critical GAIN-7TM interaction motifs unique to each compact state and investigated their roles in signal transduction through single-point mutagenesis (**Methods, Fig. 4C-I**). When reaching the TA-coupled state, L10 first partially uncouples from the α-domain (as observed in the isolated GAIN WT_LP_ state) to reach an intermediate state stabilized by polar interactions between ECL2 (Arg 541) and GAIN (Ser 363, Cys 366, Glu 367) (**Fig. 4C**). This transient state is followed by the locking of L1 onto the 7TM through a salt bridge between Lys 195 and Asp 540 and the formation of a hydrophobic contact network between the TA (Leu 388) and ECL2 (Ile 546, Ile 547) (**Fig. 4C**). The TA-ECL2 interface is further stabilized by the formation of an extended TA-ECL2 beta-strand hydrogen-bond pairing and a polar edge between L10 (Gln 345) and ECL2 (Tyr 558) (**Fig. 4C**). In the TA-uncoupled state however, L10 first uncouples from the α-domain to reach a fully opened conformation and contact ECL2 through a salt bridge between Asp 352 and Arg 551 (**Fig. 4C**). In this intermediate state, GAIN undergoes a rigid body translation over the 7TM moving L7 and TA away from the GAIN-7TM interface. L1 then folds back onto the 7TM, creating a strong polar contact network between L1 (e.g. Lys 195) and ECL2 (e.g. Asp 566) that further stabilizes this TA-excluded interface (**Fig. 4C**). Overall, since the aforementioned hydrophobic and polar contacts are unique to each state, we replaced the corresponding residues with either Ala or Phe to shift the GAIN-7TM conformation towards the TA-uncoupled (“Shift TA uncoupled”) or the TA-coupled state (“Shift TA coupled”) (**Fig. 4D-F**).

We evaluated the impact of these mutations on ADGRG1’s constitutive activation of the Gα13-RhoA pathway. This was measured using a serum response element (SRE) luciferase assay in HEK293 cells transfected with ADGRG1 variants bearing these GAIN domain mutations (**Methods, Extended Data Fig. 15**)^31, 49, 50^. The mutants predicted to shift the GAIN-7TM conformation displayed a wide range of effects consistent with our predicted GAIN-7TM binding modes and their implication in receptor constitutive signaling (**Fig. 4E,F, Supplementary Discussion**). Mutations of the hydrophobic core led to either decreased or increased constitutive activity, strongly correlating with their impact on the packing of the interface (**Fig. 4E,F**). Interestingly, enhanced binding interactions were mediated by a movement of GAIN toward the 7TM, reminiscent of a push mechanism of activation where compression of the GAIN-7TM interface enables enhanced binding surface area conducive to signal transmission. These results validate the importance of the TA-ECL2 hydrophobic core in signal transduction and suggest that native interactions have evolved to maintain a suboptimal hydrophobic communication interface in the absence of ligand and mechanical stimuli. Overall, these findings support the involvement of specific hydrophobic and polar contacts in stabilizing the two compact states and validate our structural models of the GAIN-7TM binding interfaces, as well as the existence of an equilibrium between signaling-incompetent and signaling-competent ligand-free states of ADGRG1 (**Fig. 4G**).

### Collagen 3 binding activates ADGRG1 through conformational selection of signaling states

If this conformational equilibrium model is correct, agonist binding should be capable of activating the receptor even in the absence of mechanical stimulus through a classic conformational selection mechanism. To validate this hypothesis, we measured the impact of Collagen 3 (Col3) binding on the activation of our library of ADGRG1 variants (**Methods**). Since Col3, a known ADGRG1 agonist, binds to the PLL domain, we can rule out any direct effect of Col3 on the GAIN-7TM interface. Therefore, Col3 binding is expected to influence receptor activity primarily through long-range allosteric interactions and the stabilization of specific receptor conformations.

While Col3 binding significantly increased the activity of the WT and neutral mutants, we observed much smaller effects in the shifted variants (**Fig. 5A**). When interpreted using a Boltzmann distribution of states, these results align well with a conformational selection model (**Fig. 5B**). Specifically, if the signaling-incompetent state is too stable, Col3 binding cannot shift the equilibrium sufficiently to significantly populate the active state. Conversely, if the equilibrium is already dominated by the signaling-competent state, receptor activity reachable without any applied mechanical force is near its maximum possible value, and agonist binding has little additional impact.

**Figure 5.**
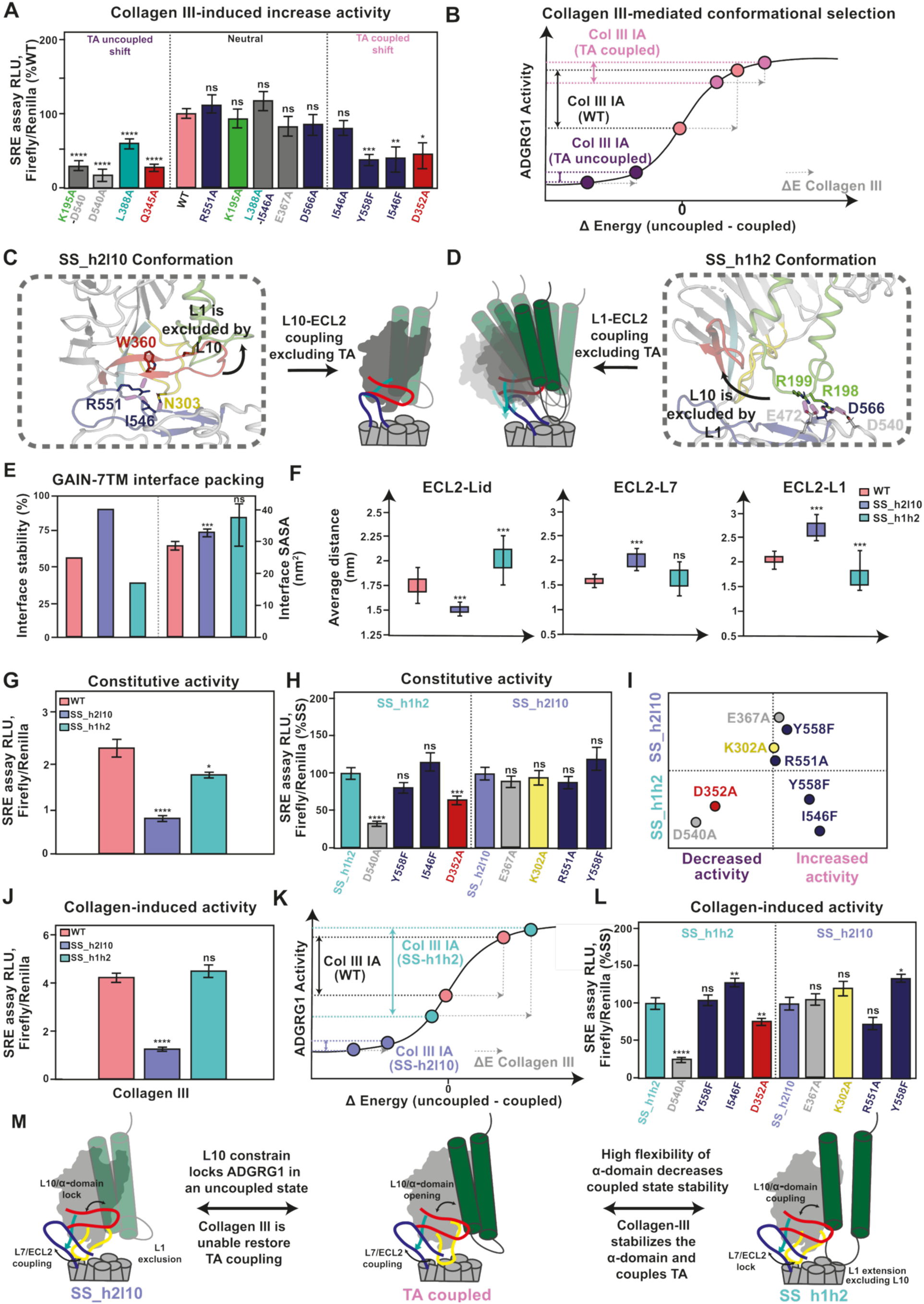
Collagen 3 stabilizes the signaling competent state through conformational selection. **A.** Impact of point mutations on the Collagen III–induced increase in ADGRG1 activity, relative to constitutive activity, as measured by SRE-luciferase assay and normalized to WT. Significant differences with respect to WT were calculated by one-way ANOVA (***: p-value < 0.001; ns: non-significant). **B.** Schematic representation of the Collagen III-induced changes in the Boltzmann distribution of uncoupled and coupled states. The y axis describes the receptor activity which is directly related to the fraction of receptors occupying the TA-coupled state. The x-axis describes the difference in energy between TA-coupled and TA-uncoupled states. The grey dotted arrow describes the stabilization of the TA-coupled state through conformational selection by collagen 3 binding (ΔE Collagen III). Changes in activity (IA) for WT, TA-uncoupled and TA-coupled shifted receptor variants upon collagen 3 binding are described as vertical black, purple and pink double arrows, respectively. **C,D.** Representative structures and schematic representations of SS_h2l10 (**C**) and SS_h1h2 (**D**) dominant conformations. **E.** GAIN-7TM interface stability (**Methods**) and variations in Solvent accessible surface area (SASA) observed in the simulations. Values for WT, SS_h2l10 and SS_h1h2 are shown in salmon, blue and green, respectively. **F.** Average distances between GAIN flexible regions and ECL2 in the simulations. Significant differences from WT were calculated by one-way ANOVA (***: p-value < 0.001; ns: non-significant). **G,J.** Constitutive (**G**) or Collagen III-induced (**J**) activity of WT and engineered ADGRG1 variants, measured by SRE-luciferase assays. Significant differences from WT were calculated by one-way ANOVA (***: p-value < 0.001; ns: non-significant). **I.** Expected impact of point mutations on the engineered variants activity. **H,L.** Impact of point mutations on the Constitutive activity (**H**) and Collagen III-induced activity increase (**L**) of the engineered variants measured by SRE-luciferase assay and normalized to the respective SS_h1h2 and SS_h2l10 variants. Significant differences with respect to WT were calculated by one-way ANOVA (***: p-value < 0.001; ns: non-significant). **K.** Schematic representation of the Collagen III-induced changes in the Boltzmann distribution of uncoupled and coupled states. The y axis describes the receptor activity which is directly related to the fraction of receptors occupying the TA-coupled state. The x-axis describes the difference in energy between TA-coupled and TA-uncoupled states. The grey dotted arrow describes the stabilization of the TA-coupled state through conformational selection by collagen 3 binding (ΔE Collagen III). The lower GAIN-7TM interface stability in SS_h1h2 results in a larger impact of collagen 3 binding (ΔE) in this scheme. Changes in activity (IA) for WT, SS-h2l10, SS-h1h2 receptor variants upon collagen 3 binding are described as vertical black, blue and green double arrows, respectively. **M.** Schematic descriptions of the designed GAIN domains’ impact on the GAIN/7TM interface, compared to the one observed for the WT ADGRG1 TA-coupled state.

### Designed GAIN domains lock ADGRG1 into signaling incompetent conformations

Since the motions of helices H1, H2, and loops L1, L7, and L10 are critical for the formation of the signaling-competent state, we next examined how the reprogrammed dynamic behaviors of these switchable elements in the designed GAIN domains impacted ADGRG1 basal and agonist-induced activities. We began by conducting long equilibrium MD simulations of ADGRG1 variants bearing the designed GAIN domains SS_h1h2 and SS_h2l10, referred to as ADGRG1_h1h2 and ADGRG1_h2l10, respectively. Our observations revealed significant effects of these designs on the orientation of the extracellular region and the interaction of GAIN and TA with the 7TM domain (**Fig. 5C-F**).

In the ADGRG1_h2l10 variant, where L10 is constrained to H2 and unable to undergo the opening seen in the GAIN WT_LP_ and ADGRG1 TA-coupled states, the receptor should be unable to access the signaling-competent state, a prediction confirmed by our simulations and AlloPool analysis. Instead, the GAIN-7TM interface adopted a single, highly stable conformation, dominated by polar contacts between L7, L10, and ECL2, which excluded L1 and TA from the interface. Consistent with this stable TA-excluded state, ADGRG1_h2l10 exhibited no significant basal or Col3-induced activity in the absence of mechanical stress (**Fig. 5H–J**). Furthermore, single-point mutations failed to disrupt the interface and restore basal signaling (**Fig**. **5H, L**).

In our simulations of ADGRG1_h1h2, we observed a greater diversity of ECR conformations compared to the WT, due to increased flexibility of the GAIN/PLL domains. Unlike the WT, where H1 coupled with PLL while H2 primarily connected to L7 and L10, the engineered disulfide bond locked both helices to the PLL, transforming the GAIN-PLL subdomains into a rigid body. As a result, the GAIN domain lost its ability to dynamically adapt its conformation to the 7TM, leading to decreased packing and destabilization of the GAIN/7TM interface (**Fig. 5D-F**). In most observed receptor structures (**Fig. 5D**), L1 contacted ECL2 through an extended conformation (similar to WT TA uncoupled state), which excluded L10 and TA from the GAIN/7TM interface.

Consequently, ADGRG1_h1h2 was expected to occupy the TA-coupled state less frequently than the WT, consistent with its lower basal activity and the reduced effect of point mutations on signaling activity signaling (**Fig. 5H, L**). However, due to the weakened GAIN-7TM interface, Col3 binding had the largest impact among all variants, triggering more than a twofold increase in ADGRG1_h1h2 activity (**Fig. 5H-L**).

Overall, our findings in the absence of mechanical stimulus support a receptor activation model driven by an equilibrium between signaling-incompetent and signaling-competent conformations of the GAIN/7TM interface. The transition to the signaling state is controlled by the precise orientation of the extracellular region and the coordinated motions of key GAIN switchable motifs, which result in partial TA exposure and coupling to the 7TM’s extracellular loop 2. As demonstrated by our engineered receptors, perturbations in GAIN loop and alpha-domain dynamics can significantly influence access to signaling states and the regulation of receptor activity by agonists through a conformational selection mechanism (**Fig. 5M**).

### Shear stress direction dictates shedding-dependent versus independent mechanotransduction

We next investigated how the application of mechanical force impacted the GAIN-7TM interface conformations and receptor signaling. Instead of attempting to replicate the most physiologically relevant but complex mechanical cues sensed by ADGRG1 in native tissues^13^, we designed a simple *in vitro* assay that probes cellular responses upon adhesion to a solid surface coated with collagen III, a known adhesion ligand of ADGRG1. In this assay, cells expressing ADGRG1 receptors were subjected to vibration-induced mechanical stimulation (**Method**). While this setup may not recapitulate the physical state of the ligands in the extracellular matrix, it allowed us to directly compare the signaling of cells expressing distinct ADGRG1 variants under variable and quantitatively reproducible mechanical stimuli, and to assess the impact of the designed molecular perturbations of the GAIN. Vibration treatments enhanced the signaling responses of all the ADGRG1-expressing cells with the largest increases measured for the SS_h1h2 followed by the WT (**Fig. 6A**).

**Figure 6.**
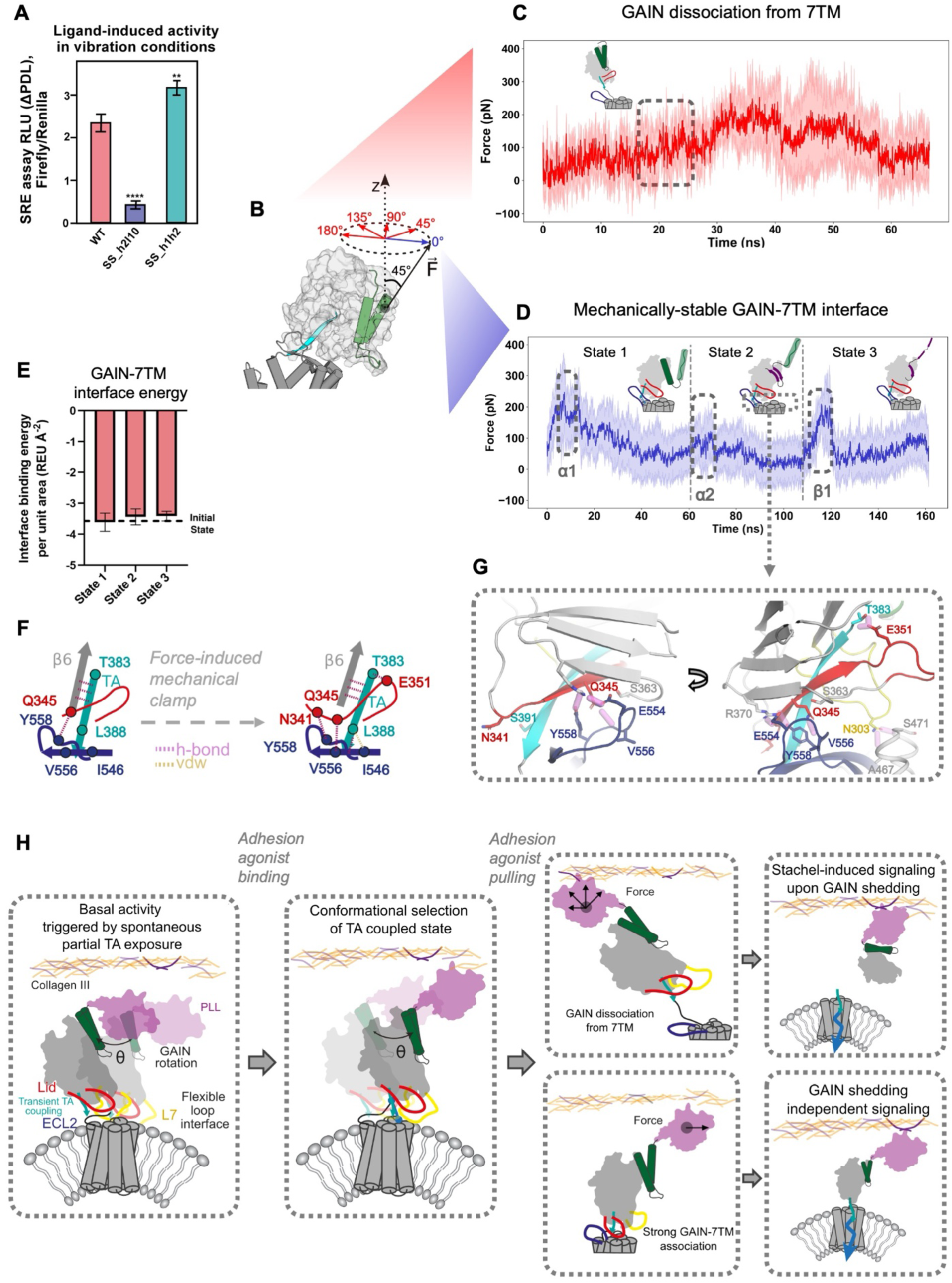
Shear stress direction dictates shedding-dependent versus independent mechanotransduction. **A.** Collagen III-induced activity in vibration conditions, measured by SRE assays. Significant differences compared to WT were calculated by t-test (**: p-value < 0.001, ****: p-value < 0.0001). **B.** Schematic representations of 5 pulling directions along which the force is applied during SMD. Red-colored directions lead to GAIN dissociation from the 7TM. Blue-colored direction maintains strong GAIN-7TM association. **C.** Representative force-extension curve of the dissociation. **D.** Representative force-extension curve maintaining strong GAIN-7TM association. Boxed areas indicate unfolding events: helix 1, helix 2 and beta-strands 1&2. **E.** GAIN-7TM interface energy along the SMD trajectories maintaining GAIN-7TM association. **F.** Schematic representation of the force-induced mechanical clamp formation at the GAIN-7TM interface involving enhanced interactions between TA, L10 and ECL2. Hydrogen bonds and Van der Waals contacts are represented as pink and yellow dotted lines, respectively. **G.** Structural snapshots of GAIN-7TM interactions under mechanical load. Hydrogen bonds are represented as pink lines, respectively. **H.** Proposed mechanisms of ADGRG1 constitutive, ligand-induced, and force-induced signaling.

To better understand the structural origin of these effects, we ran SMD simulations on the WT receptor. In the absence of structural information on the binding mode of adhesion ligands to the receptor ECR, we reasoned that applied shear stress would exert pulling forces on the ECR along different directions depending on the ligand’s binding orientation. To model these possibilities, we performed SMD simulations by pulling the ECR along five uniformly distributed directions within the x–y plane parallel to the cell membrane (**Fig. 6B**). In all cases, the applied force rapidly triggered the dissociation of GAIN’s helices from the GAIN’s beta sheets, driving the domain into a fully loaded conformation. Interestingly, this conformation resembles that of the ADGRG1_h1h2 variant in the absence of mechanical input. This suggests that GAIN SS_h1h2 permanently occupies a loaded state conformation, implying a higher susceptibility to mechanical activation than WT, consistent with its largest increase in signaling response to mechanical stimulus.

Subsequently, we observed rapid dissociation of the GAIN domain from the 7TM surface in four of the five pulling directions (**Fig. 6C**), whereas in one direction, the GAIN remained stably associated with the transmembrane region (**Fig. 6D-G**). In this latter case, we detected mechanically induced conformational rearrangements at the GAIN–7TM interface—primarily involving L10 and ECL2—that resulted in the formation of “mechanical clamps”. These clamps reinforced contacts between the TA (Leu 360) and L10 (Asn 313, Gln 317) with ECL2 (Glu 383, Val 528, Tyr 530), and between L7 (Asn 275) and the extracellular tip of TM 2 (Ala 439, Ser 443) (**Fig. 6G**). In this conformation, the TA was further stabilized at the interface compared to the TA-coupled state observed without applied force (**Fig. 6F**). Strikingly, these interactions remained intact even under forces sufficient to unfold the GAIN alpha-domain and the first beta strands (**Fig. 6D,E**). These results indicate that, in this particular direction, the GAIN–7TM interface is mechanically robust, suggesting a shedding-independent mode of mechanotransduction. Conversely, in the other pulling directions, GAIN rapidly dissociated from the 7TM, eliminating critical interactions required for allosteric signaling between the ECR and 7TM. Under these conditions, mechanotransduction would likely rely on shedding-dependent TA release from the GAIN.

## Conclusion

Our study provides an unprecedented molecular blueprint of ADGRG1 activation, elucidating how the GAIN domain mediates critical communication between adhesion and signaling regions (**Fig. 6H**). In our model, GAIN coordinates receptor signaling by integrating diverse chemical and mechanical cues and converting them into specific conformational states of its α-subdomain, modulated loop dynamics, and TA exposure. These features collectively shape the ECR–7TM interface at the cell surface. In the absence of external stimuli, constitutive activity arises allosterically from large-scale ECR fluctuations that occasionally sample a signaling-competent conformation, marked by partial TA exposure and transient interaction with ECL2. Ligand binding enhances this activity, likely by constraining ECR motion and selectively stabilizing the signaling-competent state. Under mechanical load, two distinct modes of mechanotransduction emerge depending on the direction of applied force: GAIN either dissociates from the 7TM domain to promote signaling via shedding, or remains engaged with the 7TM to drive activation through allosteric coupling.

Although shedding-dependent and -independent mechanisms have often been viewed as mutually exclusive in aGPCR signaling, our findings suggest that a single receptor can toggle between these pathways depending on the nature of the mechanical input. As aGPCRs frequently bind multiple adhesion ligands, some of which adopt distinct physical forms, the ligand-bound extracellular regions (ECRs) of a single receptor may span a wide range of structural configurations and adhesion modes. Combined with the diversity of possible shear stress directions, this implies that aGPCRs operate within a constantly shifting landscape of mechanical cues. The mechanistic versatility we uncovered may enable these receptors to decode such complex mechanical inputs into selective chemical responses within the cell.

Our findings offer a molecular framework that integrates both allosteric and mechanical perspectives on aGPCR activation, facilitating the exploration of aGPCR functional diversity and aiding drug discovery efforts. Additionally, our study validates a powerful computational approach for understanding and designing protein motions and mechanics, areas that remain largely untapped in biomolecular engineering.

## Acknowledgments

We thank Demet Araç for sharing the dualLUC-SRE plasmid, Luciano Abriata for help in setting up the CD experiments, the Barth lab members and Alex Persat for insightful discussions and critical comments on the manuscript.

## Author contributions

PB, MM and LD conceived the project. PB supervised the entire project. MN supervised the AFM measurements. FP supervised the protein production and the X-ray crystallography. LD made constructs, purified protein, performed and analyzed CD and cell signaling assays. LD and MM carried out the protein modeling and design. MM developed the force propagation pathway analysis and design approaches. MM, MP and MH performed the SMD calculations. BSY carried out and analyzed the AFM measurements. KL, LD and ANL produced the proteins in insect cells. KL, LD and ANL carried out the protein crystallization. FP and KL oversaw the X-ray data collection and structure determination. PB and MM wrote the manuscript. All authors provided comments on the manuscript.

## Data and materials availability

The X-ray structure data are deposited in the Protein Data Bank under the pdb code 9S9U. The AFM and SMD data are available upon request. All software codes developed in this study are available at https://github.com/barth-lab/AlloPool. All other data are available in the main text or the supplementary materials.

## Extended Data

**Extended Data Figure 1.**
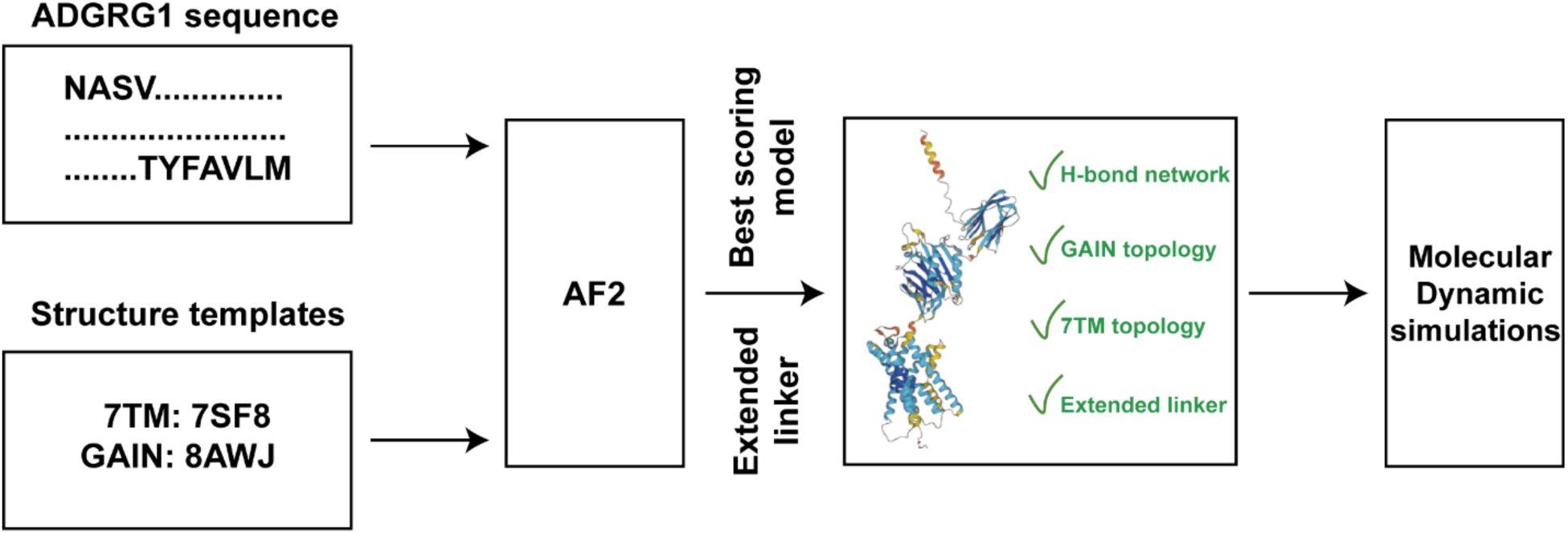
Computational pipeline for modeling ADGRG1 structure and dynamic. Alpha-Fold (AF2) was used to predict the structure of ADGRG1 starting from 2 different templates: 7SF8 (active state of the TM domain) and 8AWJ (structure of the human GAIN domain of ADGRG1). The best model was selected based on different criteria such as 1) the conservation of the hydrogen-bond network stabilizing the TA within the GAIN domain, 2) the conservation of the GAIN topology, and 3) an extended GAIN-7TM linker conformation. Molecular Dynamic simulations were performed on the best model.

**Extended Data Figure 2.**
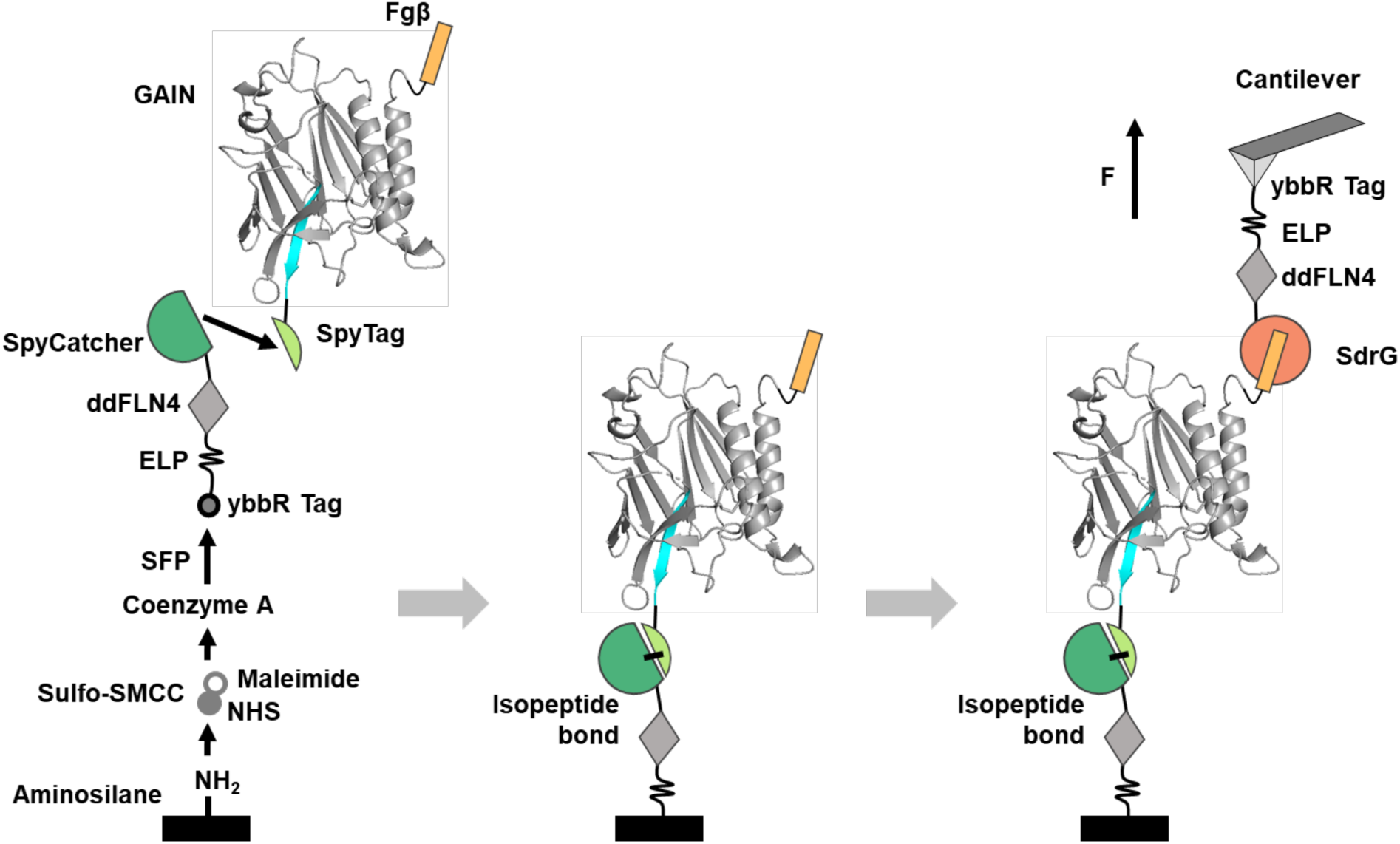
Schematic of the GAIN construct designed for the AFM-SMFS experiments. Schematic steps of the cantilever and coverglass surface modification and protein immobilization for AFM-SMFS experiments (**Methods**).

**Extended Data Figure 3.**
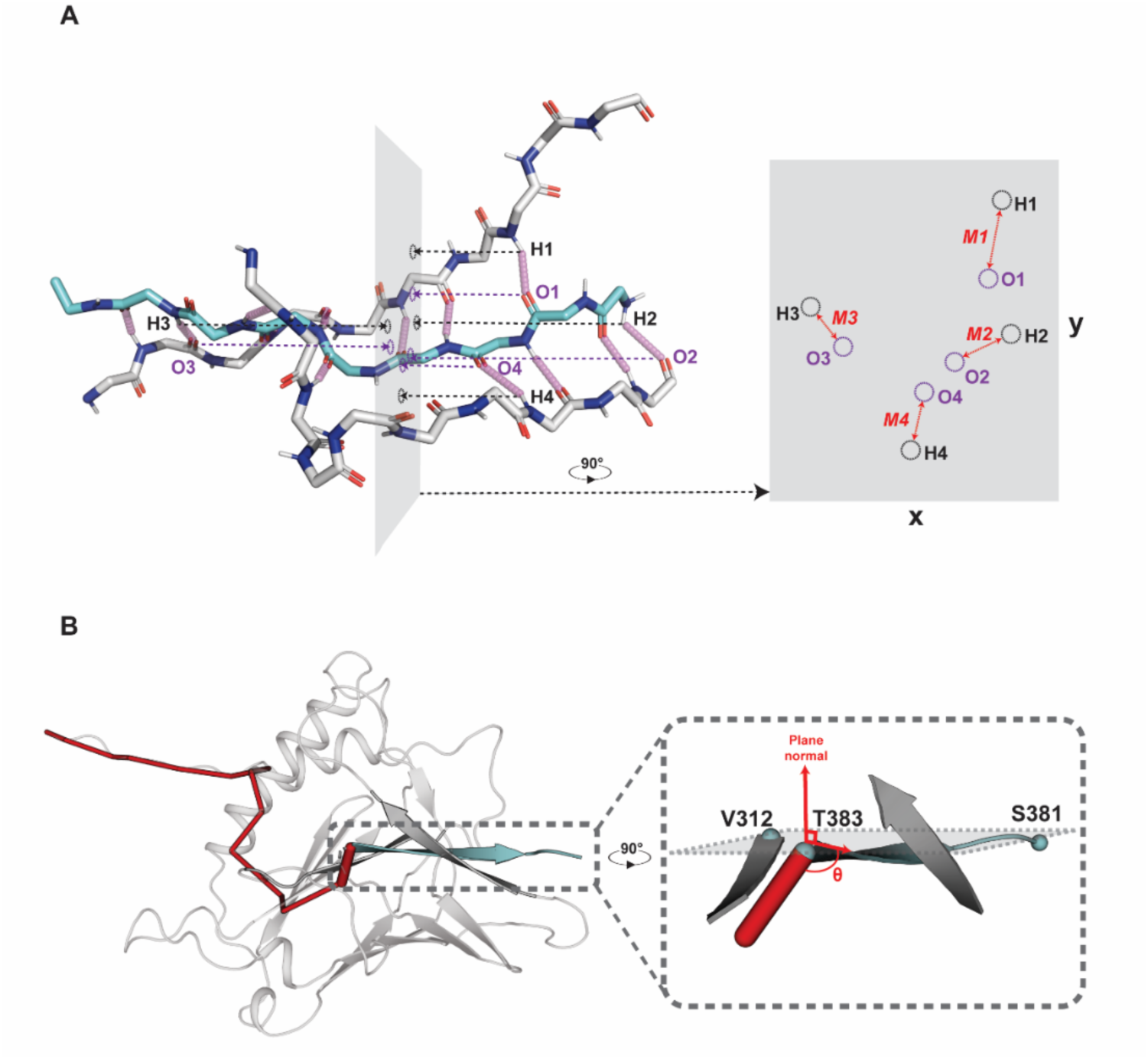
Schematics of the calculation methods used for force propagation angle and H-bonds magnitude. **A.** H-bonds are projected on a 2D plane perpendicular to the pulling axis (pointing on the C-terminus of the TA) and their magnitude are computed within this plane. Rotation of the plane shows the different magnitudes (M) that are computed for each hydrogen bond and compared between the different states. **B.** The WT GAIN structure in the loaded state and the corresponding force propagation pathways (red tubes). Right: zoom on the TA (cyan) and β-strands 6 and 8 (gray). The force (red tube) reaches the TA on T383. The path angle (Theta) is computed with respect to a vector lying in the plane defined by the 3 annotated residues and orthogonal to the plane normal (**Methods**).

**Extended Data Figure 4.**
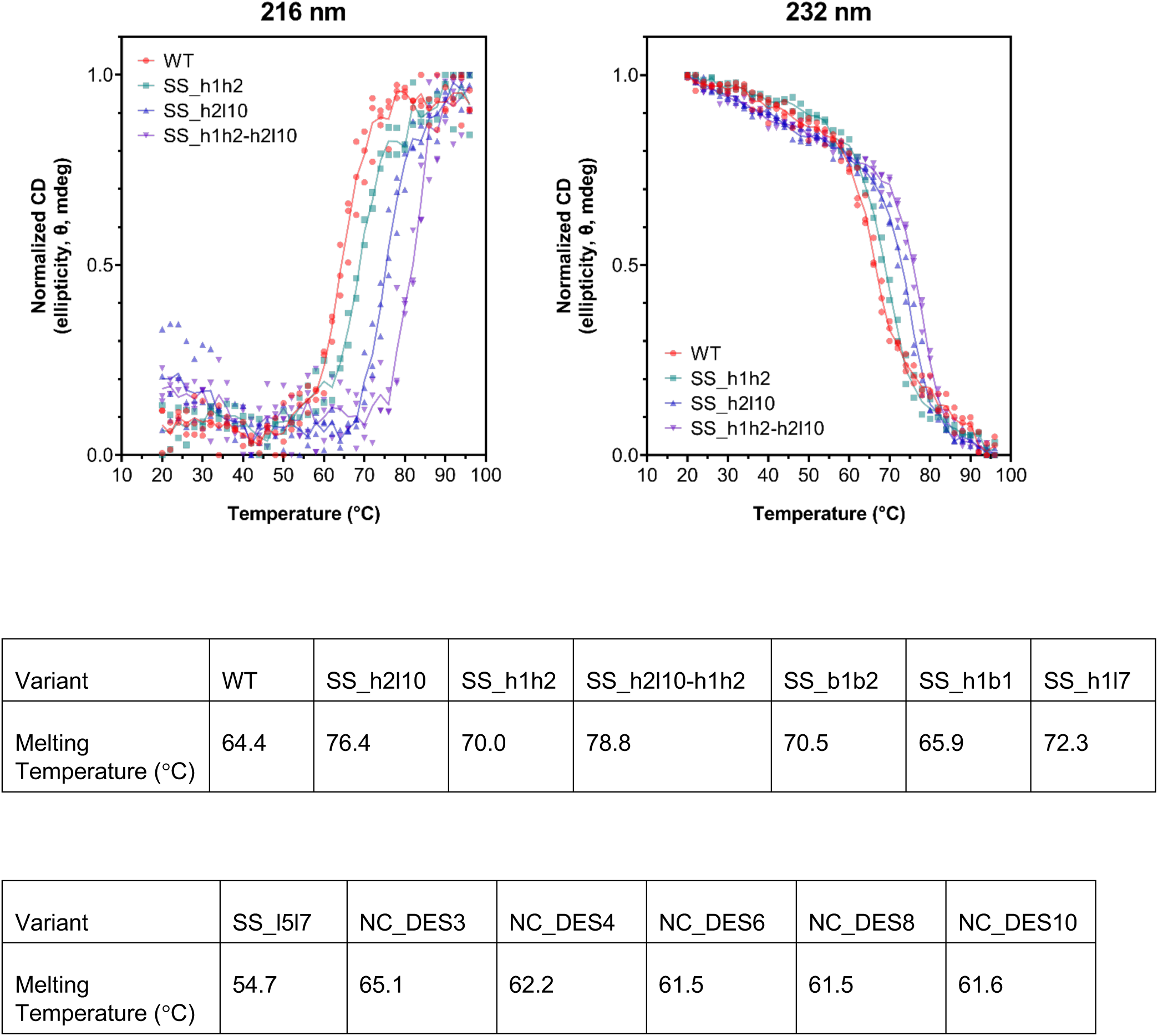
Thermal melts of the GAIN domains measured by Circular Dichroism. Thermodynamic stability of purified GAIN domain variants measured by circular dichroism spectroscopy (CD). CD melting curves from 20°C to 96°C at 216 nm and 232 nm are shown for WT and three GAIN domain disulfide variants. Melting temperatures are reported in the tables for WT, variants designed with disulfide bridges (denoted SS_*) and variants designed through non-covalent interactions (denoted NC_DES*).

**Extended Data Figure 5.**
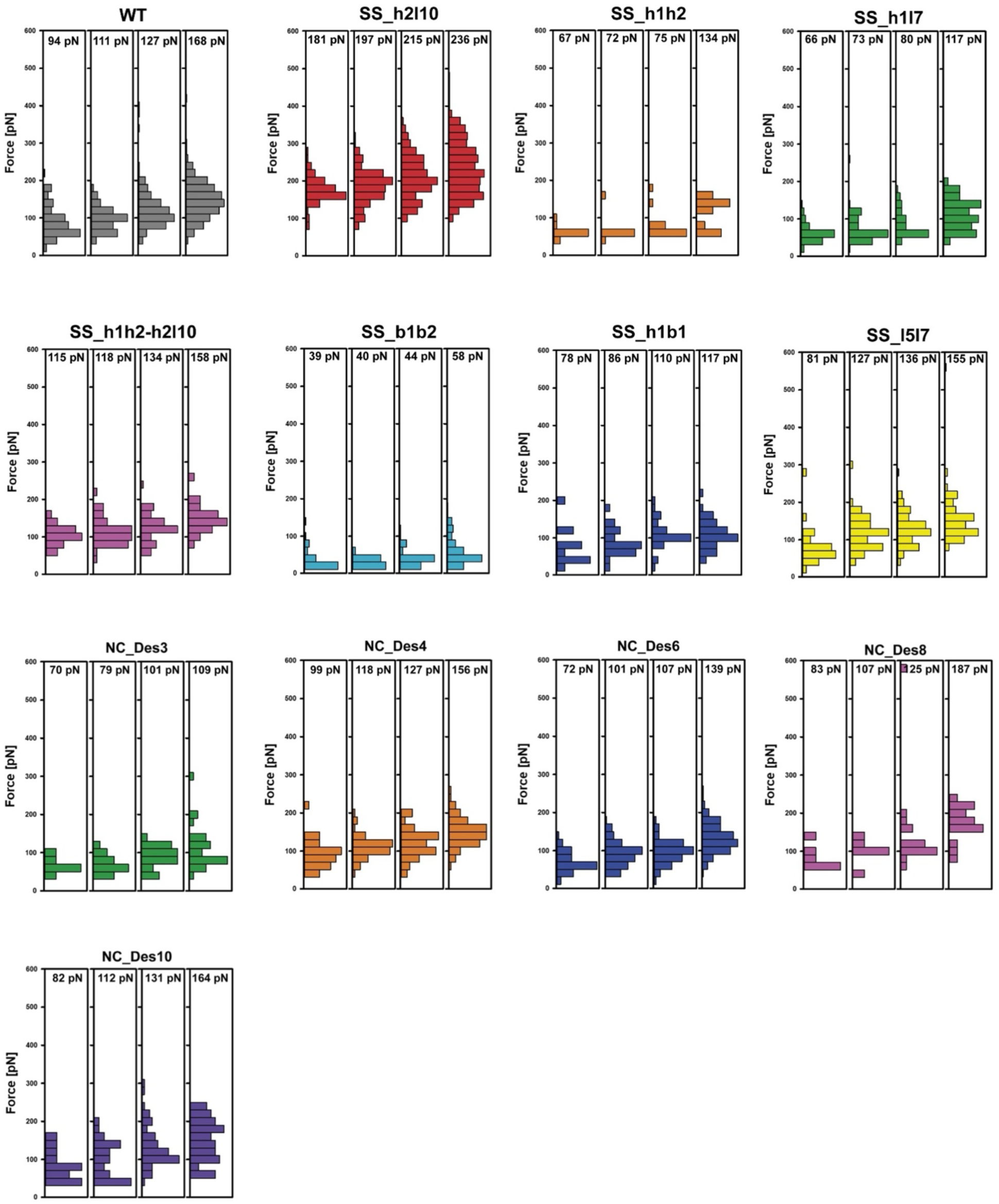
GAIN-TA Rupture force analysis by AFM. Distribution and median values of GAIN-TA dissociation forces recorded for WT and designed variants of GAIN at four distinct pulling speeds (from left to right): 400 nm/s, 800 nm/s, 1600 nm/s, 3200 nm/s.

**Extended Data Figure 6.**
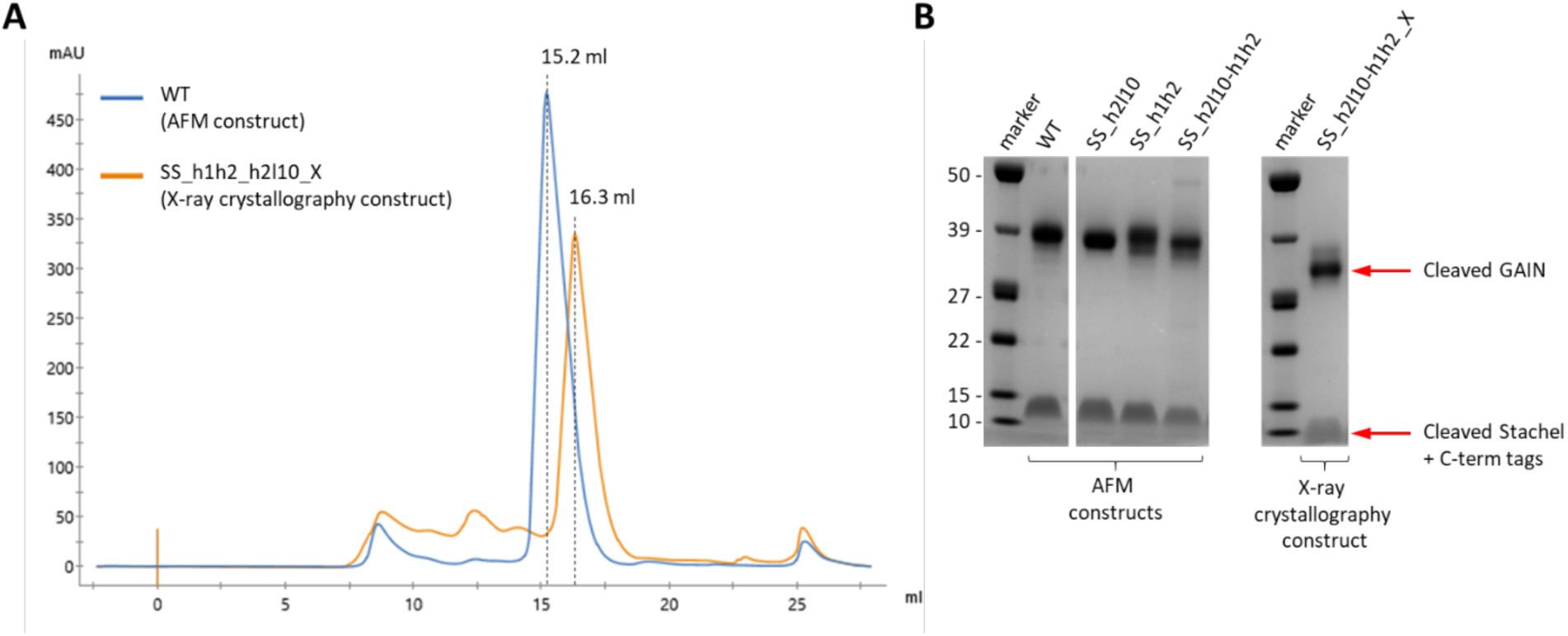
GAIN domain purification for AFM and X-ray crystallography. **A.** Gel-filtration (Superdex 200 Increase 10/300 GL) elution profiles for representative purifications of the WT GAIN domain construct for AFM experiments and of the SS_h1h2-h2l10 GAIN domain construct for X-ray crystallography (suffix “X”). Protein constructs for X-ray crystallography lack the N- and C-terminal AFM handles (see full protein sequences). **B.** SDS-PAGE analysis of the purified WT and variant GAIN domains and of the SS_h1h2-h2l10_X GAIN domain construct. The lower molecular weight bands correspond to the cleaved C-terminal containing the TA and tags and/or handles.

**Extended Data Figure 7.**
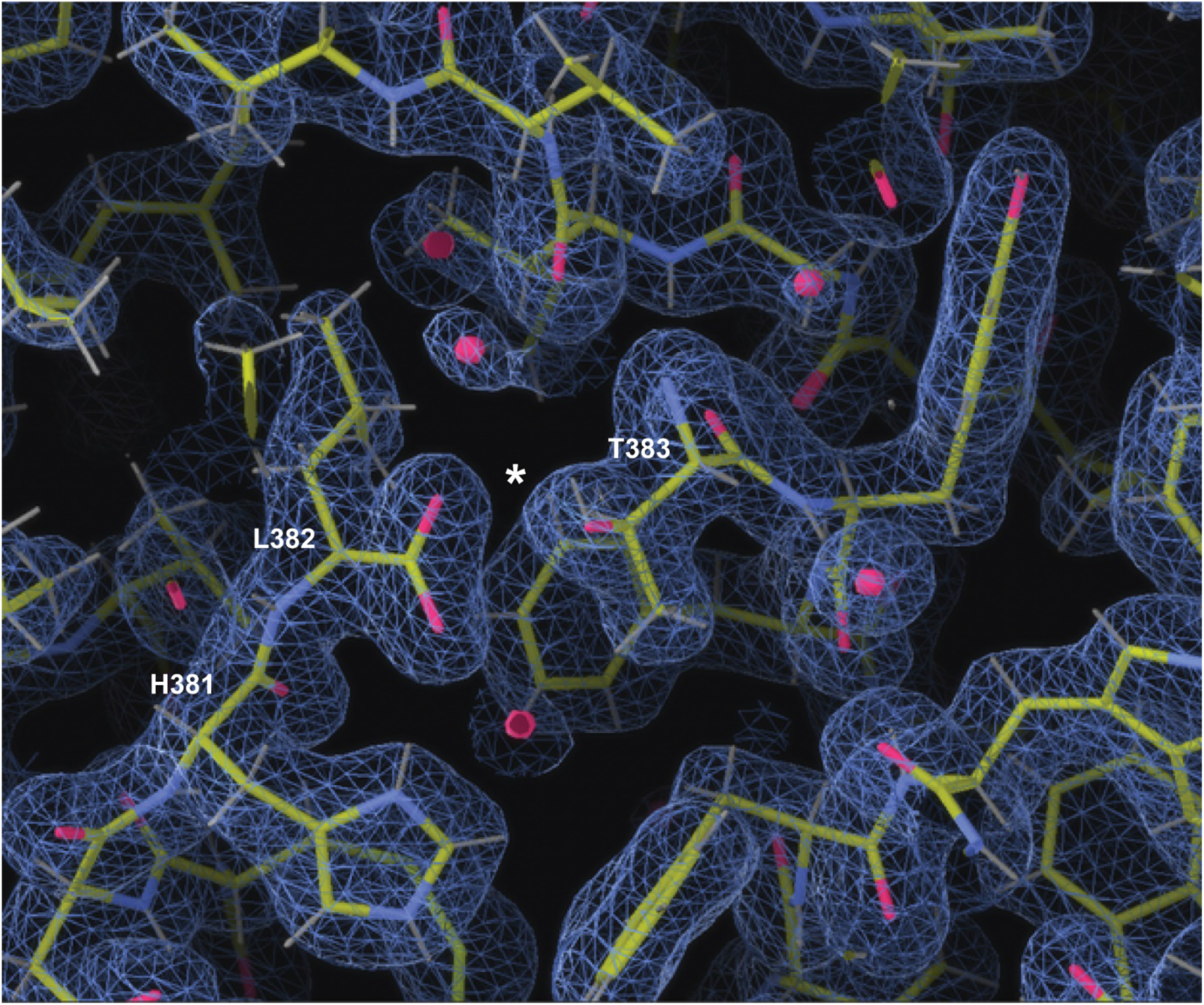
Electron density map of the GAIN domain near the autoproteolytic site. Electron density (2Fo-Fc) map contoured at 1 sigma at the C-terminus of the designed GAIN domain (L382) and the N-terminus of the TA peptide (T383). The cleavage site is represented by a white star.

**Extended Data Figure 8.**
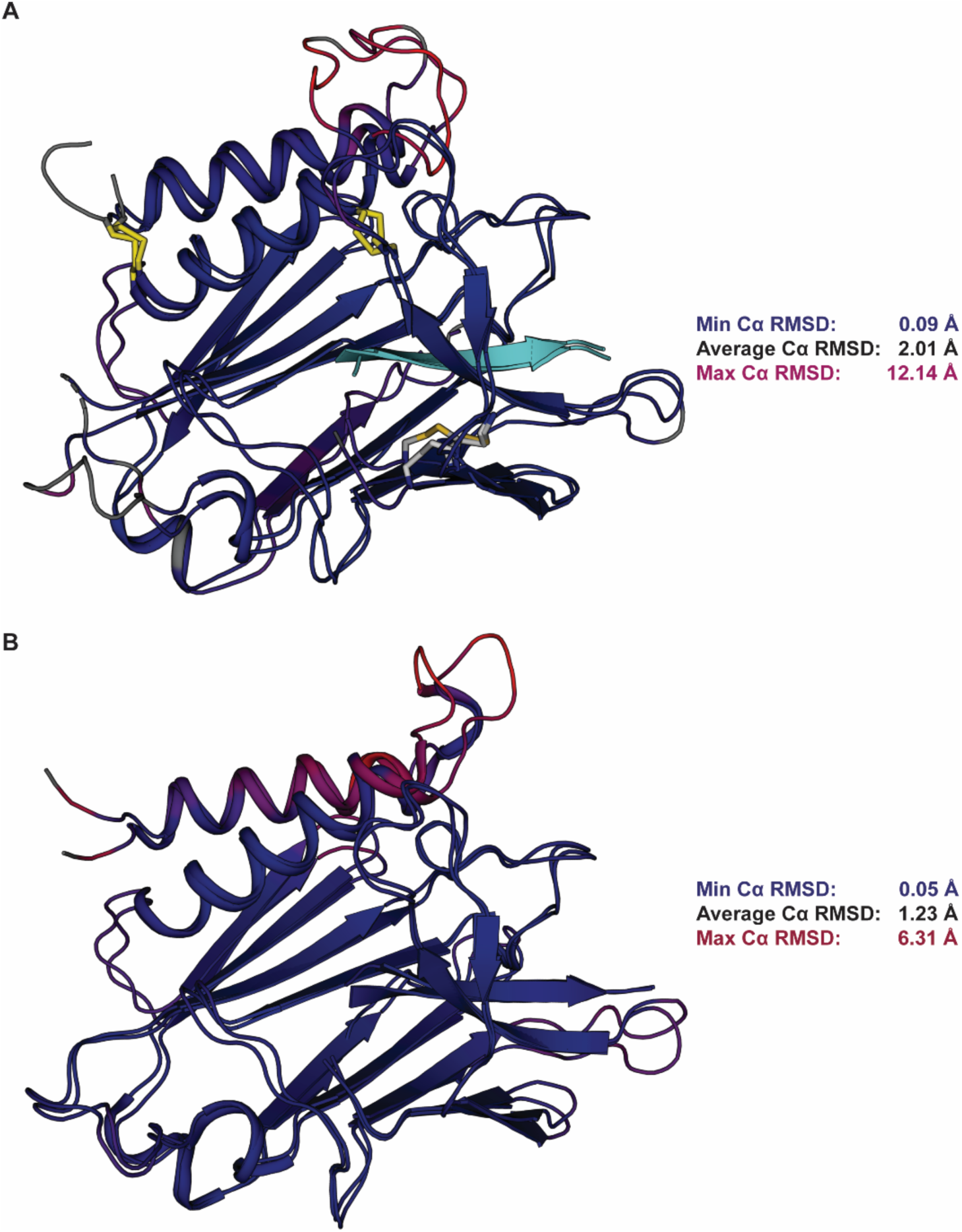
Comparison between the experimentally-determined and predicted GAIN structures. **A.** Structural alignment of the *in-silico* predicted SS_h1h2-h2l10 mutant and its crystal structure colored by *α*-carbon RMSD. **B.** Structural alignment of the crystal GAIN domain with the murine structure (5KVM) colored by alpha carbon RMSD.

**Extended Data Figure 9.**
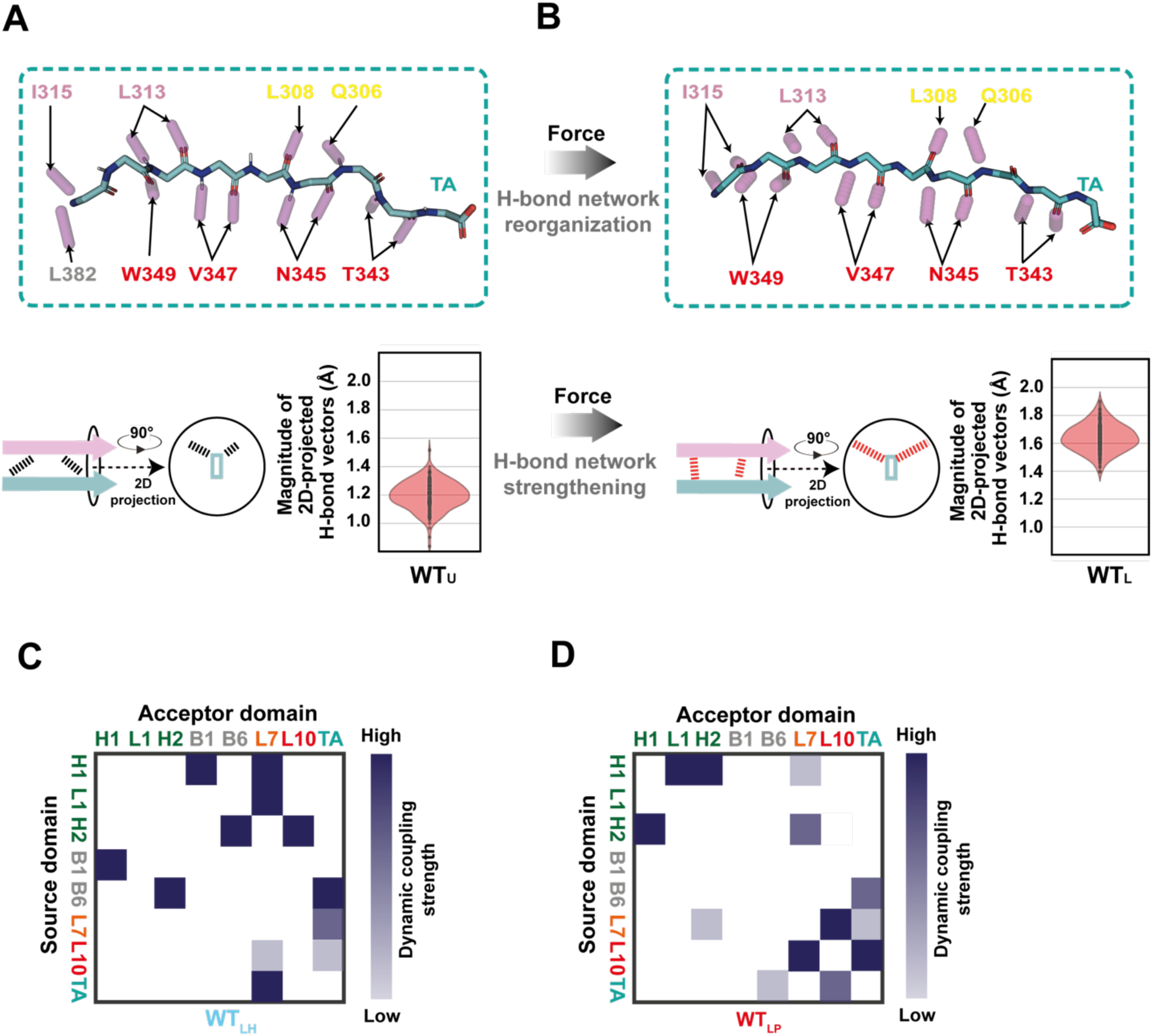
Impact of mechanical loading on GAIN WT. **A, B.** Reorganization of the hydrogen bond network between TA and GAIN upon mechanical load. **A.** Hydrogen bond network in the resting state. Structural representation of the TA with individual hbonds (pink sticks) to neighboring residues of the GAIN (above). Magnitude of backbone hydrogen bond vectors projected on a plane orthogonal to the direction of the pulling axis (below). Larger magnitudes of H-bond 2D projections correspond to geometries with larger orthogonal components to the pulling force axis, and can be interpreted as a stiffer and more mechanically resistant H-bond interface (**Methods**). **B.** Hydrogen bond network in the mechanically loaded WT_LP_ state. Structural representation of the TA with individual hbonds (pink sticks) to neighboring residues of the GAIN (above). 2D projection of the hbond network (below). **C, D.** 2D coupling maps between local regions of the GAIN domains. Coupling values are derived from the pooling edge frequencies calculated by AlloPool in the mechanically-loaded WT_LH_ (**C**) and WT_LP_ (**D**) states (**Methods**). The direction of the information propagation from source to acceptor domains corresponds to the direction of the pooled edges (**Methods**). The relative strength of the dynamic coupling ranges from low (light gray) to high (dark grey).

**Extended Data Figure 10.**
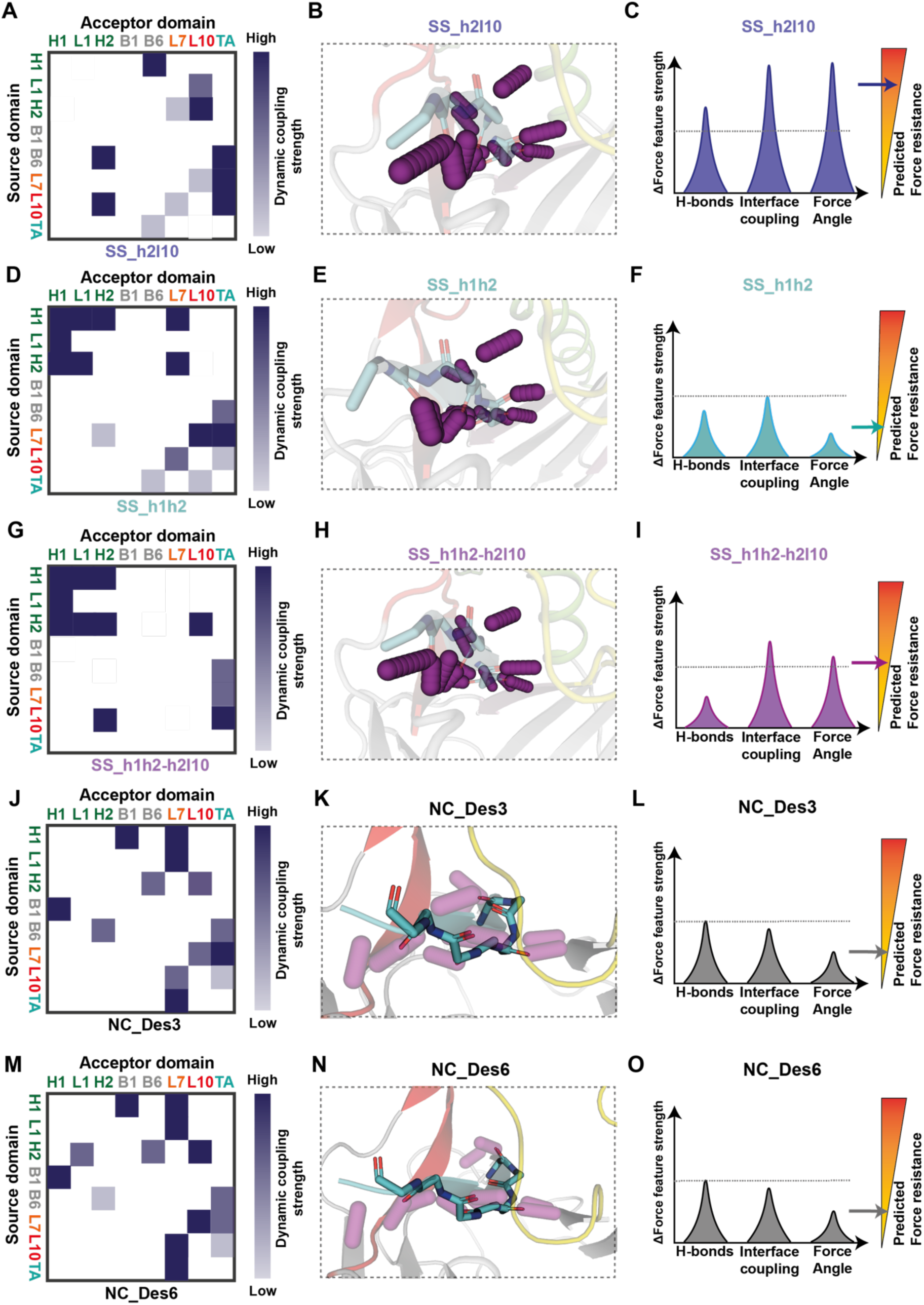
Mechanical features of GAIN variants. **A, D, G, J, M.** 2D coupling maps between local regions of the GAIN domains. Coupling values are derived from the pooling edge frequencies calculated by AlloPool in the mechanically-loaded states (**Methods**). The direction of the information propagation from source to acceptor domains corresponds to the direction of the pooled edges (**Methods**). The relative strength of the dynamic coupling ranges from low (light gray) to high (dark grey). **B, E, H, K, N.** Backbone H-bond networks between the TA and neighboring beta-strands and loop L7. Individual hydrogen bonds are represented as magenta sticks. **C, F, I, L, O.** Schematic description of the mechanical feature strength and the resulting predicted mechanical resistance of the GAIN-TA interface (indicated by an arrow) for each variant compared to WT (dotted line). Force angle features are described in **Extended Data Figure 12**.

**Extended Data Figure 11.**
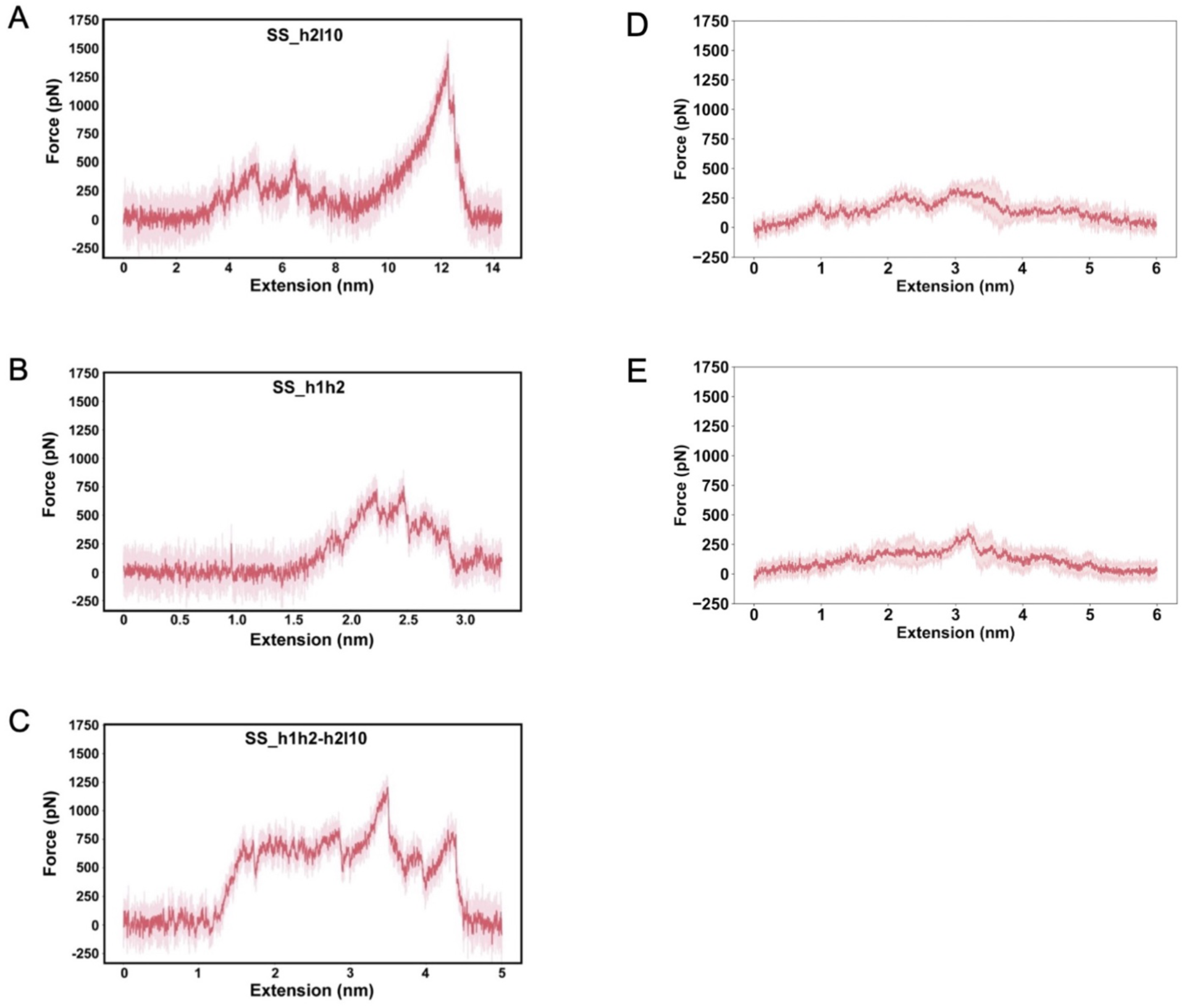
Representative SMD curves of the highest force resistance peaks for designed GAINs. Force versus extension curves for the SS_h2l10 (**A**), SS_h1h2 (**B**), SS_h1h2-h2l10 (**C**), NC_Des3 (**D**) and NC_Des6 (**E**) designs. Simulations were performed at a pulling speed of 0.1 nm.ns^−1^ (**Methods**).

**Extended Data Figure 12.**
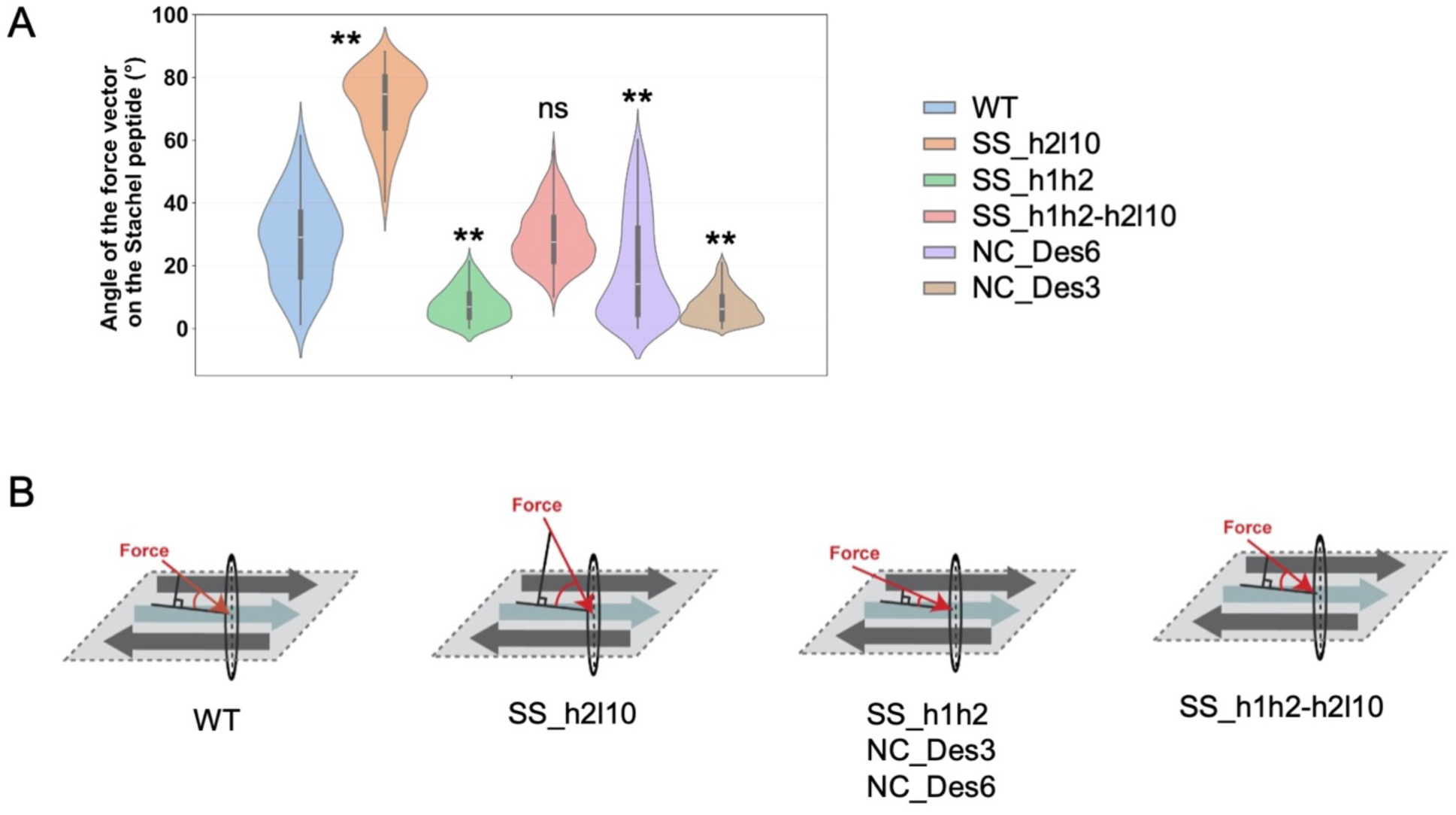
Force propagation pathway projection in the engineered variants. **A.** Angle of force vector on the TA peptide for WT and engineered GAIN domains (ns: non-significant; * p-value < 0.05; ** p-value < 0.01 as compared to WT) at 0.1 nm.ns^−1^ pulling speed. **B.** From left to right: schematic representation of the major force propagation pathway projection in the fully loaded states of the WT, SS_h2l10, SS_h1h2, SS_h1h2-h2l10, NC_Des3 and NC_Des6 variants.

**Extended Data Figure 13.**
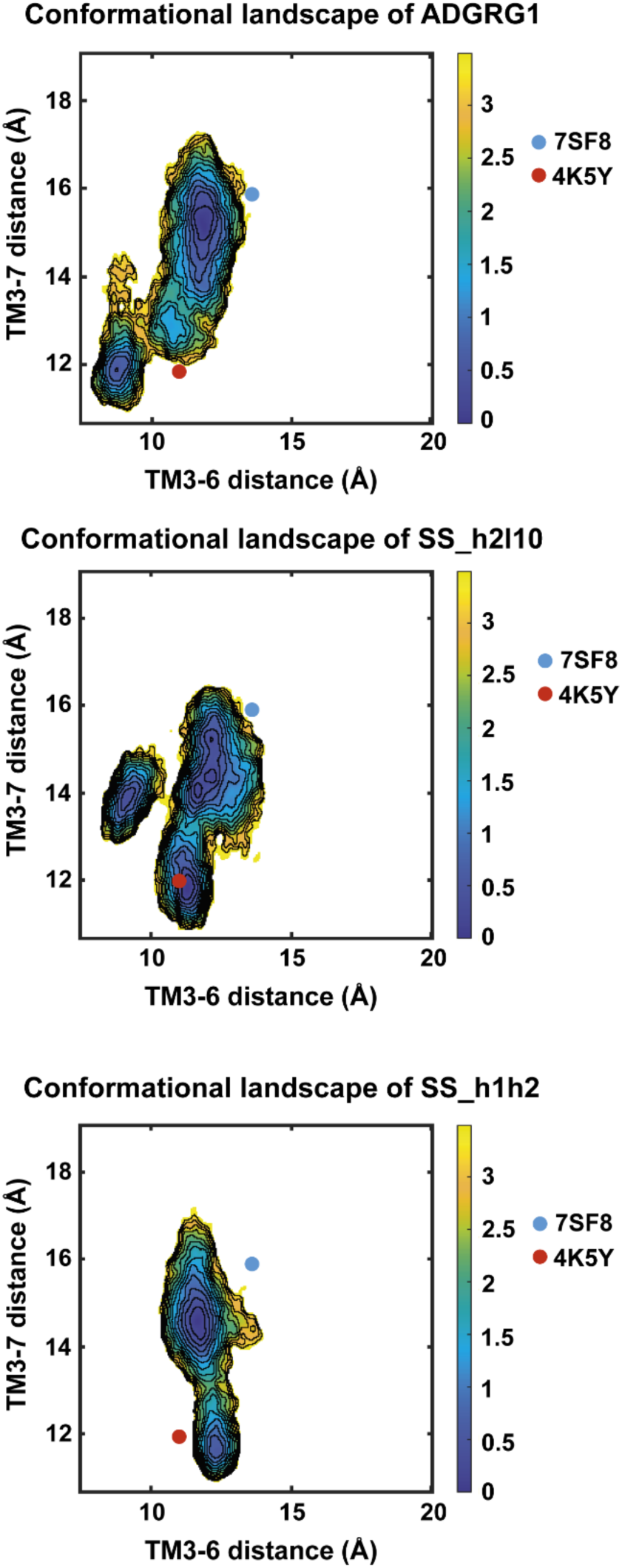
Conformational landscape of ADGRG1 and engineered variants. Conformational landscape analysis extracted from the Molecular Dynamic simulations of WT (top), SS_h2l10 (middle) and SS_h1h2 (bottom) ADGRG1 variants. Residues used for TM3-6 and TM3-7 distances are 3.50, 6.42 and 7.56 (in Wootten numbering). References for active state of ADGRG1 (PDB: 7SF8) and inactive state of a class-B GPCR (PDB: 4K5Y, corticotropin-releasing factor receptor 1) are indicated for comparison.

**Extended Data Figure 14.**
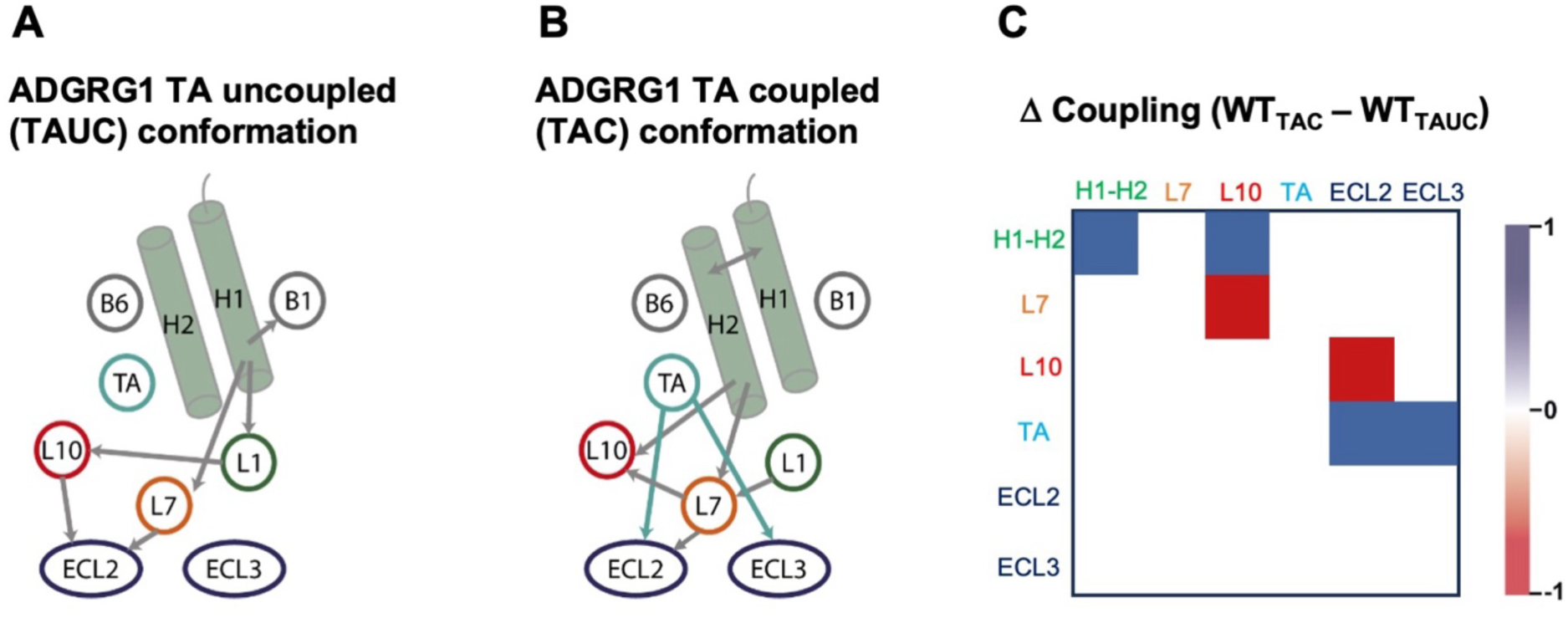
TA-mediated allosteric communication in WT ADGRG1. **A, B**. Schematic representation of the subdomain interaction graph in ADGRG1 for the two observed compact conformations of WT ADGRG1. Directed information propagations are represented by arrows. **C.** Difference of subdomain coupling between the TA-coupled (TAC) and the TA-uncoupled (TAUC) conformations. Stronger allosteric coupling of the TA with ECL2 and ECL3 is observed in the TA-coupled conformation.

**Extended Data Figure 15.**
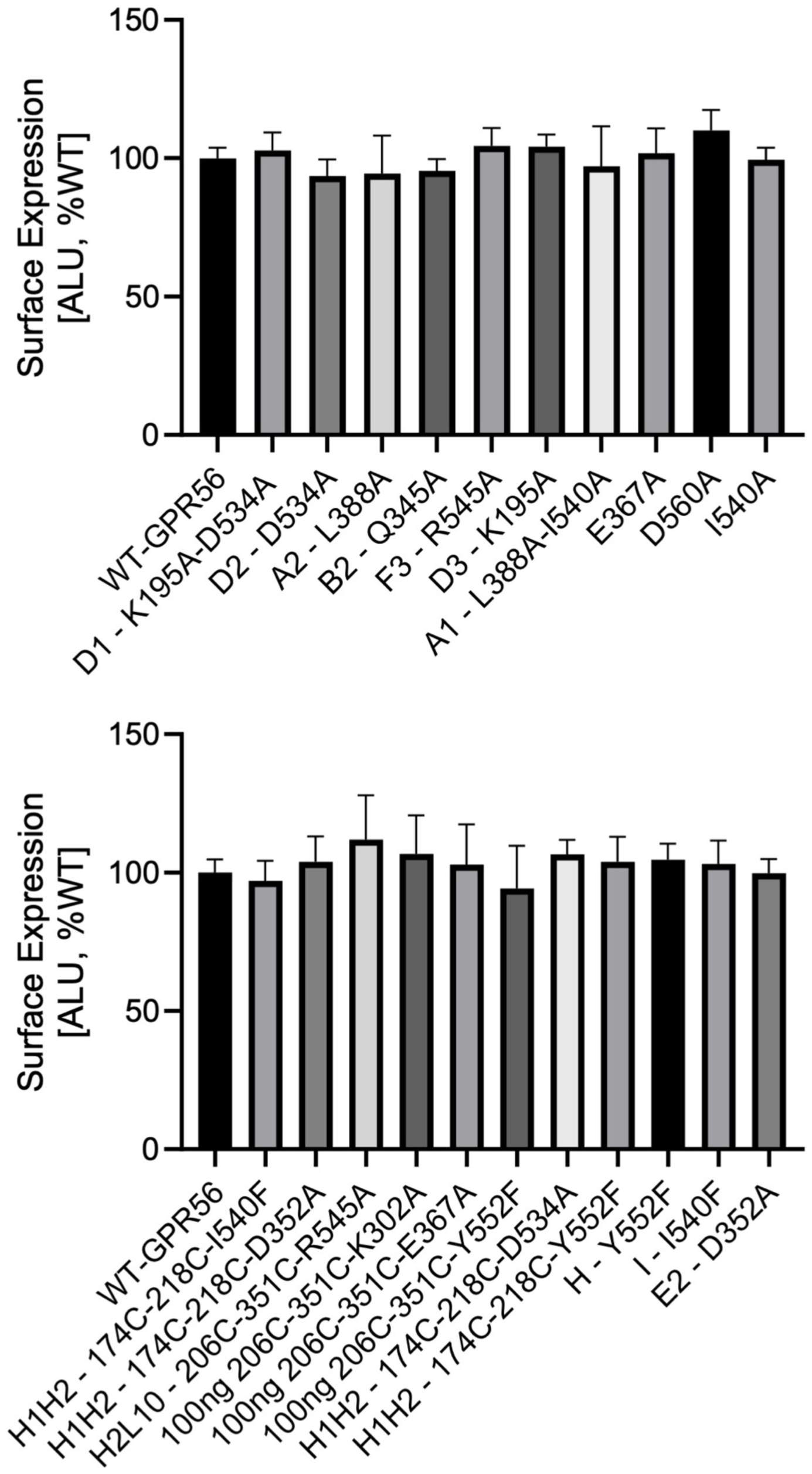
Cell surface expression of ADGRG1 variants. The receptor surface expression was quantified using ELISA (**Methods**). The values reported are the mean and standard deviations of 3 independent experiments.

**Extended Data Table 1.**
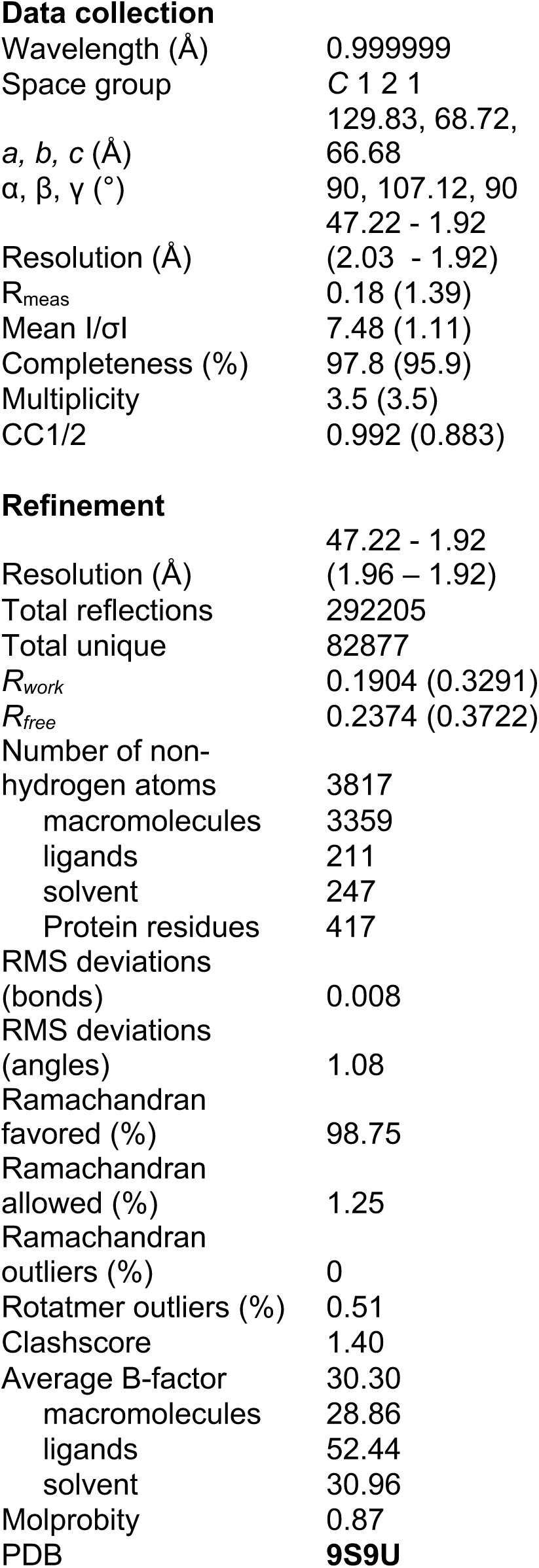
X-ray crystallography, molecular replacement and refinement data.

## Supplementary Information

### Supplementary Discussion

#### Structural characterization of the GAIN SS_h1h2_h2l10 variant

To ensure that the engineered disulfide bridges were correctly folded, we characterized the designed GAIN structures using X-ray crystallography (**Methods, Extended Data Figure 6, Extended Data Figure 7**). While the murine ADGRG1 GAIN structure was solved in the presence of the ligand adhesion PLL domain and a bound antibody^31^, we obtained crystals diffracting at 1.9 Å for the human SS_h1h2-h2l10 GAIN construct alone (**Extended Data Table 1)**. The structure revealed an overall β-sandwich and α-domain structure similar to that of the murine form (Cα RMSD = 1.2 Å), but diverged significantly in the loop regions due to low sequence and structure homology in these regions (**Extended Data Figure 8**). The structure matched our homology models very closely and the conformation of the designed disulfide bridges were superimposable to our predicted bonds (Cα RMSD = 2 Å, **Extended Data Figure 8**).

#### Mechanical stability analysis of disulfide-locked GAIN designs

**SS_h2l10.** The newly-identified loaded state of this variant characterized by the formation of a Y-lock motif exhibits several features reminiscent of highly mechanically stable interfaces (**Extended Data Figure 10A-C**). Compared to WT (**Extended Data Figure 9**), we observed a redistribution of the polar contacts and an increased helicoidal geometry of the hydrogen backbone network engaged by the TA, which is reminiscent of highly mechanically stable interfaces (e.g. sdrG-Tgb, ^26^) (**Extended Data Figure 10B**). Unlike in WT where only one β-strand and L10 are dynamically coupled to TA (**Extended Data Figure 9**), all components of the TA-GAIN interface are strongly coupled in the SS_h2l10 variant (**Extended Data Figure 10A**), ensuring an optimal distribution of the mechanical force over the entire interface, thereby enhancing its mechanical resistance, similarly to what was observed in the SdrG-FgB interface^26^.

**SS_h1h2.** Our analysis identified additional features that were predictive of a mechanically looser interface in the SS_h1h2 variant (**Extended Data Figure 10D-F**). While the distribution of hydrogen bonds within the rupture interface was identical to WT (**Extended Data Figure 9A**), we observed an increased planarization at the polar network and overall loss of coupling between the TA and the rest of the GAIN domain, which should weaken the interface (**Extended Data Figure 10D-F**). These features validate the design rational and indicate that the h1h2 variant cannot reach more mechanically stable loaded states (**Extended Data Figure 10F**).

**SS_h1h2_h2l10.** In our simulations, the variant first behaves similarly to SS_h1h2 before reaching a higher mechanically resistant state stabilized by a partial Y-lock motif (**Fig.3K**). The presence of the two disulfide locks causes extreme stretching of the alpha-domain, leading to the decoupling of L7 and L10 (**Extended Data Figure 10G**). This prevents the formation of an optimal Y-lock interaction network, making it less mechanically resistant compared to SS_h2l10. Further accumulation of force within GAIN disrupts this loaded state, leading directly to a two-step dissociation of the TA through an unzipping-like mechanism. While the helicoidal geometry of the backbone polar network around the TA was reminiscent of h2l10, the variant lost 2 hydrogen bonds with L7 from the TA at the origin of the unzipping-like TA release (**Extended Data Figure 10H**). The coupling between neighbor β-strands and TA was also weakened, suggesting a less resistant interface than in SS_h2l10 (**Extended Data Figure 10G,I**).

#### Mutagenesis of polar contacts at the WT GAIN-7TM interface

To validate our predicted conformations of the WT GAIN-7TM interface and their relevance for signal transduction, we identified critical GAIN-7TM interaction motifs unique to each conformational state and investigated their roles in signal transduction through single-point mutagenesis. We replaced the corresponding residues with either Ala or Phe to shift the GAIN-7TM conformation towards the TA-uncoupled (“Shift TA uncoupled”) or the TA-coupled state (“Shift TA coupled”) (**Fig. 4D-F**). The mutants predicted to shift the GAIN-7TM conformation displayed a wide range of effects consistent with our predicted GAIN-7TM binding modes and their implication in receptor constitutive signaling (**Fig. 4E,F**).

Specifically, with the exception of the Asp352Ala mutation, substitutions of polar residues were either functionally neutral or led to a loss of function. For example, mutations at Arg551 and Lys195 had no significant impact on constitutive activity, consistent with their involvement in polar interactions in both the signaling-incompetent (TA-uncoupled) and signaling-competent (TA-coupled) compact states. In the TA-coupled state, Arg551 forms polar contacts with multiple residues (Ser363, Cys366, and Glu367). Thus, disrupting a single interaction—such as via the Glu367Ala mutation—is insufficient to shift the equilibrium between the two states, explaining the lack of functional effect of that point mutation. Similarly, in the TA-uncoupled conformation, Lys195 interacts with Asp566, but this interaction can be preserved through compensatory contacts with neighboring acidic residues (e.g., Asp538) upon the Asp566Ala substitution. This adaptability accounts for the neutral effect of the Asp566Ala mutation on receptor activity.

However, mutation of Asp 540, which interacts with Lys 195 exclusively in the TA-coupled state, was expected to specifically destabilize that state. Consistent with this prediction, the mutation led to a significant decrease in ADGRG1 constitutive activity. Conversely, the mutation of Asp 352 enhanced constitutive activity, in line with the destabilization of the TA-uncoupled conformation and a shift toward the signaling-competent state.

Overall, these findings support the involvement of specific polar contacts in stabilizing the two compact states and validate our structural models of the GAIN-7TM binding interfaces, as well as the existence of an equilibrium between signaling-incompetent and signaling-competent ligand-free states of ADGRG1.

## Materials and Methods

### AlphaFold model of ADGRG1

The structural model of ADGRG1 was obtained using AlphaFold (*51*). We used the sequence of the human ADGRG1 (uniprot ID: Q9Y653) along with two structural templates for the TM domain (PDB: 7SF8) and the GAIN domain (PDB: 8AWJ) to predict multiple structures of ADGRG1. We filtered the modeled structures using the pLDDT score and the following metrics: (i) conservation of the canonical backbone hydrogen bond network stabilizing the TA in the GAIN domain, (ii) accuracy of the predicted GAIN topology compared to the crystal structure (PDB: 8AWJ), (iii) accuracy of the predicted TM domain compared to referenced active state structure (PDB: 7SF8), (iv) extended linker between the TM-domain and the TA to explore a large conformational space of the GAIN-7TM putative interactions.

### Design of GAIN domain constructs for SMFS-AFM experiments

We designed the GAIN domain constructs for AFM experiments based on the following rationale: It is known that a protein’s mechanical response is highly dependent on the loading point and direction of the applied force (*28*, *37*–*41*). In the context of the full-length ADGRG1 receptor expressed at the cell surface, mechanical force is likely transduced to GAIN by the N-terminal adhesion PLL domain bound to an ECM ligand. To best mimic this native orientation and direction of the applied force in our SMFS experiments, we designed a setup where the C-terminal end of GAIN is attached to a solid surface through an ddFLN4 fingerprint domain while the pulling force is applied to the N-terminal end of the domain (**Fig. 1A, Extended Data Figure 1**). To enable precise repeated measurements of TA dissociation by AFM and subsequent quantitative analysis, we modified the cantilever with an SdrG-ddFLN4 fusion. This protein can reversibly bind to the Fgb peptide that was fused to the N-terminus of the GAIN constructs (**Extended Data Figure 1**). The SdrG-Fgb complex can withstand very high mechanical forces (over 2 nN) (*26*). After TA dissociation from GAIN, the cantilever is clogged with the N-terminal fragment of the GAIN. However, SdrG has a very high off-rate in the absence of force, and dissociates spontaneously from the Fgb peptide, freeing up the SdrG molecule on the cantilever for binding to new GAIN molecules on the surface. This experimental design enables precise repeated measurements and quantification of TA dissociation by AFM.

### Protein Expression and Purification

*AFM handles* – SpyCatcher and SdrG proteins for SMFS were prepared as previously reported (*28*, *52*). Constructed recombinant plasmids pET28a-ybbR-His-ELP(MV7E2)3-FLN-SpyCatcher (Addgene #157674) and pET28a-SdrG-FLN-ELP-His-ybbr (Addgene #168047) were transformed into E. coli BL21 (DE3) strain. Cells were cultured in 5 ml of Luria-Bertani (LB) medium with 50 μg ml^−1^ kanamycin at 37 °C overnight. The culture was transferred to 100 mL of Terrific Broth (TB) medium with 50 μg ml^−1^ kanamycin and cultivated at 37 °C and 200 rpm until an optical density at 600 nm (OD600) of ∼0.8-1.0 was reached. Recombinant protein expression was induced upon addition of 0.5 mM isopropyl-β-D-thio-galactopyranoside (IPTG) and the culture was further incubated at 20 °C and 200 rpm for 18 h. The cells were harvested by centrifugation at 4,000 g for 20 min at 4 °C. The cell pellets were stored at -80 °C until further purification. For purification, the harvested cell pellet was resuspended in a lysis buffer (50 mM Tris, 50 mM NaCl, 0.1% Triton X-100, 5 mM MgCl2; pH 8.0), and disrupted with a sonic dismembrator. The lysate was centrifuged at 14,000 g for 20 min at 4 °C. The supernatant was collected and incubated with Ni-NTA resin (Thermo Fisher Scientific), loaded onto a column, washed with wash buffer (1x PBS buffer with 25 mM imidazole; pH 7.2), and eluted in elution buffer (1x PBS buffer with 500 mM imidazole; pH 7.2). The eluted protein solution was buffer-exchanged and further purified by Superose SEC column, and finally stored in 33% glycerol at -20 °C.

*GAIN domain* – The ADGRG1 GAIN domain was expressed and purified following published protocols (*31*). Insect codon-optimized human ADGRG1 ECR sequences (Uniprot: Q9Y653) were synthesized (Twist Biosciences) as part of larger constructs for either AFM or X-ray crystallography experiments (see full protein sequences) in a pFastBac vector backbone with a BiP secretion signal. A baculovirus expression system was used for expression of proteins in High-Five insect cells. pFastBac constructs were transformed into DH10MultiBac cells (Geneva Biotech), white colonies indicating successful bacmid recombination were selected, and bacmids were purified by the alkaline lysis method. Sf9 cells were transfected using the X-tremeGENE 9 transfection reagent (Sigma) with the desired bacmid. eYFP-positive cells were observed after 1 week and subjected to one round of viral amplification. Amplified P2 virus was used to infect High-Five cells at a density between 1–2 × 10^6^ cells/ml. Cells were incubated for 72 h at 28°C, pelleted (1000 x *g*, 10 minutes) and the recovered supernatant was clarified (5000 x *g*, 30 minutes) and filtered using a Grade GD 1 μm pre-filter (1841-047, Whatman, GE) on top of a Durapore 0.45 μm PVDF Membrane filter (HVLP04700, Merck Milipore). 5 ml of 50% Fastback Ni Advance Resin slurry (Protein Ark) pre-equilibrated in HBS buffer containing 10 mM HEPES (pH 7.2) and 150 mM NaCl was added to the 500 ml clarified supernatant containing the secreted proteins and incubated over-night at 4°C on a rotating wheel. The resin was recovered on a gravity-flow column, equilibrated with 30 ml HBS and washed with 30 ml HBS containing 30 mM imidazole. Proteins were eluted with 15 ml of HBS containing 300 mM imidazole and 1 ml fractions were collected and analyzed by SDS-PAGE. The three or four purest and most concentrated fractions were pooled and concentrated on a 30 kDa Amicon Ultra-4 centrifugal filter (UFC803024, Merck Millipore) down to 500 μl. The concentrated proteins were loaded on a Superdex 200 Increase 10/300 GL (28-9909-44, Sigma-Aldrich) connected to an ÄKTA Pure chromatography system (GE) and eluted in HBS. The fractions containing the peak corresponding to the purified GAIN construct were pooled and quantified using A_280_ absorbance. Purified GAIN proteins were either directly concentrated to 10 mg.ml^−1^ for crystallization or flash-frozen in liquid nitrogen and stored at -80°C for subsequent AFM experiments.

### AFM-SMFS surface preparation and protein immobilization

The surface modification of cantilever and coverglasses and the protein immobilization were done as described previously (**Extended Data Figure 1**) (*52*). Cantilevers were cleaned by UV-ozone treatment for 40 min and coverglasses were soaked in piranha etching solution and rinsed with distilled water (DW). Then, cantilevers and coverglasses were treated with 3-Aminopropyl (diethoxy) methylsilane (APDMES, ABCR GmbH, Karlsruhe, Germany) to silanize the surfaces with amine groups. The amine groups subsequently reacted with an NHS group from sulfosuccinimidyl 4-(N-maleimidomethyl)cyclohexane-1-carboxylate (sulfo-SMCC; Thermo Fischer Scientific) in 50 mM HEPES buffer pH 7.5 for 30 min. The thiol group from Coenzyme A (CoA, 200 μM) reacted with a maleimide group from sulfo-SMCC in coupling buffer (50 mM sodium phosphate, 50 mM NaCl, 10 mM EDTA, pH 7.2) for 2 hrs. Finally, the ybbR-tagged proteins SdrG-FLN-ELP-His-ybbr and pre-conjugated ybbR-His-ELP(MV7E2)3-FLN-SpyCatcher-SpyTag-GAIN variants were site-specifically anchored to the surface using SFP-mediated ligation to CoA in Mg2+ supplemented 1x PBS buffer. This resulted in covalent immobilization of SdrG and GAIN variants to cantilever and cover glasses, respectively. Protein-immobilized cantilevers and coverglasses were extensively washed and kept in 1x PBS buffer prior to immediate use. SpyTag-SpyCatcher conjugation of ybbR-His-ELP-FLN-SpyCatcher and Fgβ-GAIN-His-SpyTag variants were done by mixing two proteins with same molar ratio in PBS buffer and pre-incubation for 1-12 hrs prior to ybbR tag ligation.

### AFM-SMFS measurements and data analysis

Force spectroscopy measurements with SdrG and Fgβ-fused GAIN variants were conducted in the same manner as previously illustrated using automated AFM-based SMFS (Force Robot 300, JPK Instruments). SMFS data were recorded in 1x PBS at room temperature with constant pulling speeds of 400, 800, 1600 and 3200 nm.s^−1^. A total of 258,044 force-extension curves were acquired. Force-extension curves were filtered and analyzed by a combination of software available on the AFM instrument and custom python scripts. The majority of data traces contained no interactions, non-specific interaction, or complex multiplicity of interactions. Therefore, the data traces were filtered by searching for contour length increments that matched the lengths of the fingerprint domains, FLN (≈36 nm). Theoretical contour length increment was calculated based on the equation ΔL_c_ = (0.365 nm/AA) × (# AAs in POI) - L_f_, where ΔL_c_ is expected contour length increment and L_f_ is end-to-end length of folded protein domain. For FLN, ΔL_c_ = 36.9 nm - L_f_, where L_f_ is typically < 5nm. The total number of force-extension curves matching this criterion was 509 out of 120,780, 1,058 out of 68,376, 263 out of 42,295, and 158 out of 18,447 for GAIN-WT, GAIN_SS_h2l10, GAIN_SS_h1h2, and GAIN_SS_h2l10-h1h2, respectively.

For dynamic force spectra, the rupture or unfolding forces vs. loading rate was plotted and median forces and loading rates for each pulling speed were fitted to Bell-Evans model (*49*, *50*) to estimate the effective distance to the transition state (Δx) and the intrinsic dissociation rate or unfolding rate (k_off_) in the absence of force. Data were fitted using the Dudko-Hummer-Szabo model (*55*, *56*) with a cusp-like barrier (*v* = 0.5) to estimate Δx, k_off_, and energy barrier height (ΔG).

For direct comparison with the same cantilever, both GAIN-WT and GAIN_SS_h2l10-h1h2 were immobilized on different areas of the same surface and a single cantilever was used to probe each spot using constant speed pulling at 3200 nm s-1. After 500 approach-retraction cycles at one area, the surface was moved to the other area. This cycle was done once for each GAIN-WT and GAIN_SS_h2l10-h1h2. The total number of force-extension curves matching the criterion was 69 out of 9,484, and 67 out of 4,986 for GAIN-WT and GAIN_SS_h2l10-h1h2, respectively.

For contour length analysis in reducing condition, each GAIN-WT and noGAIN was probed with the same cantilever both in non-reducing condition (0 mM DTT in 1x PBS) and reducing condition (100 mM DTT in 1x PBS). SMFS data were recorded in 1x PBS first at room temperature with constant pulling speed of 800 nm s-1, and then buffer was changed to 100 mM DTT in 1x PBS and SMFS data were recorded with the same pulling speed. The total number of force-extension curves matching the criterion was 101 out of 1,566, and 116 out of 26,097 for GAIN-WT and GAIN_SS_h2l10-h1h2, respectively.

### Circular Dichroism measurements

Purified GAIN variant proteins were adjusted to 10-15 μM final concentration in 10 mM HEPES, 15 mM NaCl, pH 7.5. Thermal denaturation CD spectra were recorded on a Chirascan CD spectrometer (AppliedPhotophysics) from 20°C to 96°C using a 2°C step, between 215 nm and 250 nm. Each melting curve was normalized between 0 and 1 across the entire spectral range and the 216 nm and 232 nm normalized spectra from 2 or 3 replicate experiments were extracted and plotted.

### X-ray crystallization and data collection

Crystals of the GAIN_SS_h2l10-h1h2 variant were grown in sitting drops using the random microseed screening (*57*). (1) 10 mg/mL of purified protein was mixed 1:1 (200 nL) with mother liquor and crystals appeared after several days at 18°C in a buffer containing 0.1 M potassium thiocyanate and 30% PEG 2000 MME. Crystals were harvested and cryoprotected in mother liquor supplemented with 25% glycerol and frozen under liquid nitrogen.

All datasets were collected at beam line PX-III of the Swiss Light Source, Villigen, Switzerland. Datasets were collected with λ = 1.00 Å. All datasets were processed with XDS. The structure was solved by molecular replacement the program Phaser, using PDB 5KVM as the initial search model. The final structures were determined after iterative rounds of model-building in COOT and refinement in phenix.refine. The model was refined to a R/Rfree of 0.196/0.241 with excellent geometry (no Ramachandran or Rotamer outliers). Model was validated with Molprobity, with a score of 1.26. Final statistics were generated as implemented in phenix.table_one (*57*–*62*).

### Mammalian Cell Assays

*Maintenance* – Human embryonic kidney cells (HEK293T, ATCC CRL-3216) were cultured in 10 cm diameter tissue culture-treated dishes in a humidified incubator (95% air and 5% CO_2_) at 37°C in growth media (DMEM, 10% heat-inactivated FBS, 1% PenStrep). Cells were passaged every 2-3 days (up to passage 25) by lifting the cells with 0.05% trypsin in DPBS.

*Transfection* – Cells were transiently (forward or reverse) transfected with plasmid constructs using the GeneJuice transfection reagent (Novagen) following the supplier’s recommended protocol (see specific conditions for each assay below). For forward transfection, 25×10^3^ cells were seeded per well of a 96-well plate and the transfection was carried out 24h later. For reverse transfection, 50×10^3^ cells were seeded per well of a 96-well plate and directly transfected.

*ELISA* – 25×10^3^ HEK293T cells were seeded per well of a 96-well plate pre-coated with 0.1 mg.ml^−1^ poly-D-lysine (PDL, Sigma). 24 hours post-seeding, cells were transfected as described previously with 100 ng of the indicated pcDNA-AVItag-GPR56 construct and 25 ng of dualLUC-SRE plasmid (generous gift from Prof. Demet Araç). 24 hours post-transfection, cells were washed with DPBS, fixed with cold 4% PFA for 10 minutes at 4°C and washed again DPBS after PFA removal. This was followed by three consecutive 60-minute incubations with (1) 2% BSA in DPBS, (2) 0.05 μg.ml^−1^ primary anti-AVI antibody (Genscript) in 2% BSA-DPBS, and (3) 0.5 μg.ml^−1^ secondary anti-rabbit antibody (Cell Signaling) in 2% BSA-DPBS. After each antibody incubation, cells were thoroughly rinsed with DPBS. The cells were then incubated in SuperSignal Chemiluminescent Substrate (ThermoScientific) at room temperature for 10 minutes. The Flexstation 3 (Molecular Devices) plate reader and SoftMax Pro 7 software (Molecular Devices) were used to measure luminescence. Assays were performed in triplicate or quadruplicate and the luminescence signals were normalized by the background signal of empty vector transfected cells.

*Dual-Glo Luciferase SRE assays* – 25×10^3^ HEK293T cells were seeded per well of a 96-well plate pre-coated with 0.1 mg.ml^−1^ PDL. 24 hours post-seeding, cells were transfected as described previously with 100 ng of the indicated pcDNA-GPR56 construct and 25 ng of dualLUC-SRE plasmid. 8-10 hours post-transfection, the culture medium was replaced with 100 μl FBS-free DMEM for over-night serum starvation. The next morning, 60 μl of the 100 μl total well volume was removed and the Dual-Glo Luciferase SRE assay (Promega) was carried out following the manufacturer’s instructions apart from a working volume of 40 μl instead of 100 μl.

*Vibration assays with coated ligands* – 96-well plates were coated overnight at 4°C with 40 μl PDL or native human collagen III (Abcam) at 1 μg.cm^−2^ in 10 mM CH_3_COOH. The next morning, 50×10^3^ cells were reverse transfected onto the pre-coated wells as described previously with 100 ng of the indicated pcDNA-GPR56 construct and 25 ng of dualLUC-SRE plasmid. 6-8 hours post-transfection, the culture medium was replaced with 100 μl FBS-free DMEM for over-night serum starvation. The next morning, cells were placed on a Titrama× 100 vibration shaker (Heidolph Instruments) and vibrated at 450 rpm over a 4-hour induction phase (2 cycles of 1 hour vibration/1 hour static). At the end of the induction phase, the Dual-Glo Luciferase SRE assay was carried out as described above.

### Computational design of GAIN variants

To identify sequence variants that modulate the mechanical stability of the GAIN domain, we first employed ProteinMPNN (*63*) and the Coupled Moves algorithm from the Rosetta suite, using the Backrub Movers protocol (*64*), on representative structures of the WT GAIN domain in its mechanically loaded state. Our rationale was that mutations stabilizing or destabilizing this conformation should directly modulate the mechanical resistance of the domain under force. To avoid introducing mutations that compromise the thermodynamic stability of GAIN in its resting (mechanically unloaded) conformation, we next evaluated the structural impact of each designed mutation on the GAIN resting state. Each mutation was introduced into the resting state model using Coupled Moves, and 100 independent structural models were generated per design. The 10% lowest-energy structures were selected, followed by full conformational relaxation using the Rosetta relax protocol to identify the global energy minimum for each variant. Any design destabilizing the resting state structure by more than 5 Rosetta Energy Units was systematically discarded.

We applied the same computational framework to search for disulfide-stabilizing residue pairs. Candidate cysteine pairs were introduced, and their sulfur–sulfur atom distances were measured across the lowest-energy models. A cutoff of 2.15 Å was used to identify disulfide-compatible pairs, and mutations not meeting this criterion were excluded from further consideration.

Finally, selected designs were subjected to steered molecular dynamics (SMD) and AlloPool analysis to validate the predicted changes in mechanical stability. These calculations confirmed the predicted shifts in mechanical resistance and the presence of key features in the mechanically loaded states—such as altered force propagation pathways, engineered catch bond motifs, and hydrogen bond networks stabilizing the tethered agonist (TA).

### SMD simulations of isolated GAIN domains

The structure of the GAIN_SS-h2l10-h1h2 variant was back-mutated using the backrub movers protocol (*64*) to recover the individual disulfide variants as well as the WT structure. The system was solvated with a box size of 80 Å along X, Y and Z dimensions with 0.1 M of *Na*^!^ and *Cl*^“^ ions. The total system size was approximatively 130.000 atoms. Simulations were performed with GROMACS 2019.4 (*65*) using the CHARMM36 forcefield (*66*) along with the TIP3 water model (*67*) in an *NPT* ensemble at 310K and 1 bar using a velocity rescaling thermostat (with a relaxation time of 0.1 *ps*) and Berendsen barostat (with a relaxation time of 1 *ps*). Equations of motion were integrated with a timestep of 2 *fs* using a leap-frog algorithm. Each system was energy minimized using a steepest descent algorithm for 5000 steps, and then equilibrated with the atoms of the protein restrained using a harmonic restraining force in two steps, first by adding the thermostat for 50000 steps and then the barostat for 500000 steps. SMD simulations were performed using the umbrella sampling method with a pulling speed ranging from 1 Å/ns to 10 Å/ns. In all simulations, the 2 alpha carbons of the N-term and C-term residues were moved harmonically in the desired direction with constant velocity. A distance cutoff of 14 Å was used for short-range non-bonded interactions, whereas electrostatic interactions were computed using the Particle-mesh Ewald method (*68*) with a cubic interpolation of power 4.

### Force propagation pathway calculation

*Correlation-based dynamical network analysis* - We carried out a correlation-based dynamical network analysis of the GAIN motions extracted from SMD to determine force propagation paths. We first calculated the correlated motions of backbone alpha-carbons (Cα) from simulation time windows corresponding to either the unloaded (equilibration without applied force) or the polar loaded state of GAIN. Force propagation paths were assigned to the network of residues connecting the N-terminus to the TA peptide that maximized the correlated motions and were predicted to most effectively transmit the force through the structure.

Specifically, correlation between alpha carbon motions were analyzed as follows. For every frame of the simulation, the [x,y,z] cartesian coordinates of every alpha carbon were extracted. Displacement of these coordinates between 2 consecutive frames were computed and represented as a vector [x_0_-x_1_, y_0_-y_1_, z_0_-z_1_]. To establish a correlation between these displacement vectors, distance correlation was used (Equation 1).

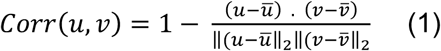

For a simulation of *t* frames, this resulted in *t* correlation matrices of [n_residues_, n_residues_]. To identify force propagation pathways, Dijkstra shortest path algorithm was used (*69*). The shortest path maximizing the total correlation were searched between the N-term of the GAIN domain and the C-term of the TA mechanical clamp for every correlation matrix, resulting in *t* shortest force propagation pathways. To extract relevant information from these pathways, k-means clustering was performed. The optimal number of clusters to use for independent simulations was determined using the inertia attribute to identify the sum of squared distances of samples to the nearest cluster center. Centroids of clusters, corresponding to the optimal force propagation pathway within this cluster were used for later extraction of features.

*Force propagation pathway projection on TA rupture interface* - To predict how force propagation paths may impact the mechanical resistance of the TA-GAIN interface, we used the angle of the force reaching that binding interface as a main criterion (**Extended Data Figure 3**). Force propagation pathways were identified as described above. To compute the angle of pathway projection on the rupture interface, Cα coordinates were extracted. These coordinates represented a [*t*,n_residues,_ n_residues_] matrix, *t* being the number of frames in the simulation. For each coordinate matrix, we constructed a plane containing both the TA N-term and C-term Cαs, and Val142 Cα. We selected Val142 (first residue of strand 6) to enable robust angle computations and avoid loop fluctuations that could introduce noise. This resulted in an ensemble of dynamic planes moving accordingly to local structural rearrangements and corresponding to the rupture interface of the TA mechanical clamp. To estimate the angle of force arrival on this rupture interface defined by these different planes, vectors connecting the last GAIN residue from where the path reaches the TA and the first TA residue reached by the path were extracted. The vectors were computed from the alpha carbon coordinates of these residue pairs. Angles between the plane normal and the force arrival vector were computed for every frame corresponding to each state of the system (unloaded, intermediate and loaded). Finally, all angles were combined from different simulation replicates, and their distribution shown as violin plots for each GAIN variant.

### Hydrogen bond 2D projection

To quantitatively assess how the backbone hydrogen bond (H-bond) network connecting the TA peptide with the neighboring GAIN beta strands responds to the applied mechanical force, we calculated the H-bond network geometry using the Wernet-Nilsson algorithm (*70*). H-bonds identified during the SMD simulations were 2D-projected onto a plane perpendicular to the TA axis defined by its N-term and C-term residues (**Extended Data Figure 3**). To highlight the H-bonds rearrangements occurring during the pulling, the magnitude of the 2D projections were compared between the native unloaded structure and the loaded structure corresponding to the highest mechanical resistance point observed in the SMD simulations. Projections were averaged across replicates and plotted. Larger magnitudes of H-bond 2D projections correspond to geometries with larger orthogonal components to the pulling force axis, and can be interpreted as a stiffer and more mechanically resistant H-bond interface.

### MD simulations of ADGRG1

The selected Alpha-Fold model for ADGRG1 was used as starting pose for MD simulations to explore the conformational space accessible to the GAIN-7TM interface. We used the Rosetta software to generate the disulfide mutants at the specific positions. The different ADGRG1 variants were inserted into a regular square POPC lipid bilayer with 170 Å side distance and solvated by a 50 Å layer of water above and below the bilayer with 0.15 M of NA^+^ and Cl^−^ ions using CHARMM-GUI bilayer builder (*71*, *72*). Simulations were performed with GROMACS 2022.4 with CHARMM36 forcefield (*73*) in a NPT ensemble at 310 K and 1 bar using a Nosé-Hoover thermostat (independently coupled to three groups: protein, membrane, and solvent with a relaxation time of 1 ps for all three) and Parrinello-Rahman barostat (with semi-isotropic coupling at a relaxation time of 5 ps respectively). Equations of motion were integrated with a timestep of 1 fs for the first three steps of equilibration and then 2 fs using the leap-frog algorithm. Each system was energy minimized using the steepest descent algorithm for 5000 steps, and then equilibrated with the atoms of the receptor and lipids restrained using a harmonic restraining force in 6 steps. After constrained equilibration, 5 independent trajectories of 400 ns each were run. The total simulated time was defined to ensure convergence of the 1^st^ and 2^nd^ order entropies calculations in the top PCA clusters in every system. While the TM-domain is in an active-state conformation in the initial structural template, the absence of the G-alpha in our simulations stabilizing the active conformation enables rapid relaxation of the TM-domain and exploration of a large space of conformations through mutual interaction with the EC region.

### SMD simulations of ADGRG1

The center of the PCA cluster representing the tethered agonist (TA)-coupled state of ADGRG1, as identified from equilibrium molecular dynamics (MD) simulations, was selected as the starting conformation for steered MD simulations. To explore the impact of the pulling directions on the conformations of the GAIN-7TM interface and accelerate the sampling, we truncated the PLL domain from ADGRG1. The structure was inserted into a regular rectangular POPC lipid bilayer with 250 Å distance perpendicular distance between parallel sides and solvated by a 150 Å layer of water above and below the bilayer with 0.15 M of NA^+^ and Cl^−^ ions using CHARMM-GUI bilayer builder. Simulations were performed with GROMACS 2022.4 with CHARMM36 forcefield in a NPT ensemble at 310 K and 1 bar with temperature coupling using velocity rescaling with a stochastic term (independently coupled to three groups: protein, membrane, and solvent with a relaxation time of 1 ps for all three) and exponential relaxation pressure coupling (with semi-isotropic coupling at a relaxation time of 5 ps respectively). Equations of motion were integrated with a timestep of 1 fs for the first three steps of equilibration and then 2 fs using the leap-frog algorithm. The system was energy minimized using the steepest descent algorithm for 5000 steps and then equilibrated with the atoms of the receptor and lipids restrained using a harmonic restraining force in 6 steps. SMD simulations were performed using the umbrella sampling method with a pulling speed of 1 Å/ns. In all simulations, the Cα of the N termini residue from the GAIN domain was move harmonically with constant velocity and in a direction specified in each round. A distance cutoff of 12 Å was used for short-range non-bonded interactions, whereas electrostatic interactions were computed using the Particle-mesh Ewald method (*68*) with a cubic interpolation of power 4. In all the simulations, we applied a harmonic constraint on the Cα of residues 437, 446, 508, 540 and 607 to prevent membrane drifting or bending.

To uniformly sample the conformational space, we considered five distinct pulling directions defined by the vectors 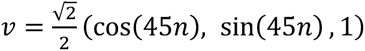 with *n* = 0, ⋯ ,4 (**Fig. 6B**). For each _-_ direction, we run four independent SMD replicates of 60 ns duration, using the same parameters as previously described. Four of the five directions led to disruption of the GAIN–7TM interface and were not pursued further. We therefore focused on the remaining direction of vector 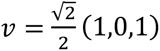, and performed 15 replicates of 160 ns each to probe the mechanical stability of the GAIN–7TM interface in detail.

During these simulations, we observed stepwise unfolding events, beginning with helix 1 (state 1) and helix 2 (state 2) of the alpha subdomain, followed by the first N-terminal strand of the GAIN domain (state 3). Protein structures were extracted at 60 ns, 110 ns, and 160 ns for further analysis. To characterize the GAIN–7TM interface, we used GetContacts to evaluate interfacial contacts and Rosetta’s InterfaceAnalyzer to compute interface energies. As most GAIN–7TM interactions remained stable between 110 ns and 160 ns, we selected from these time points the structures exhibiting the highest interface energy per unit area for detailed structural analysis presented in **Figure 6**.

### AlloPool

The AlloPool model consists of a trainable encoder-decoder architecture (*30*). The encoder predicts the interaction graph given the observed system’s dynamics, while the decoder predicts the observed trajectories given the interaction graph.

The input consists of *N* nodes representing residues in the input sequence. Each node has a feature vector describing the spatial coordinates of its C*⍺* and C*β* atoms, the backbone torsion angles *φ* and *ψ,* and a positional encoding. We denote 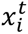 the set of features of residue *i* at time 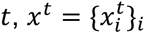. the set of all residue features at time *t*, and 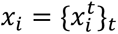 the trajectory of residue *i*. We also denote 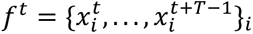. the time window starting at time *t*, where T is the number of steps. AlloPool processes each time points as a graph 𝒢_t_(*V*, *E*_t_), where *V* are nodes with associated features and *E_t_* are edges connecting nodes *(i, j)* if *d*(*i*, *j*) < 12 Å.

AlloPool simultaneously learns key edges and reconstructs the system’s dynamics in an unsupervised manner based on a learnable interaction graph *Z*(*V*, *E*_2_), where *E_p_* are pooled edges based on a learnable score associated to each edge, such that **|*E_p_*| < |*E_t_*|.** The interaction graph is then used by the decoder to reconstruct the observed trajectories.

Formally, the model minimizes the MSE between the translations 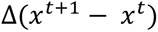 and 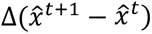, and the RMSD between *x^t^* and 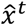 for *t* ≥ 1. We compute the RMSD using the Kabsch-Umeyama algorithm to align the structures prior to loss computation. We formalize it as 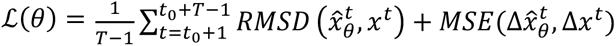 where *θ* are the trainable parameters of the model.

We leverage the node TopK operator to an Edge-TopK operator as follow:

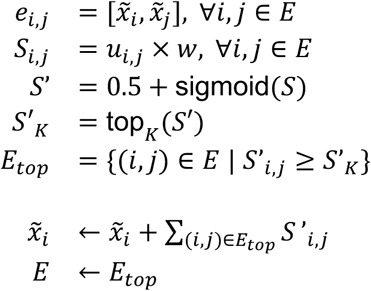

The model weights are optimized using the Adam optimizer with an initial learning rate of 10^−4^, a weight decay coefficient of 0.1, and a learning rate scheduler decreasing the learning rate by 2 after 100 optimizer steps with no loss decrease. The windows are grouped in batches of 10.

### Full recombinant GAIN protein construct sequences

The human GAIN domain sequence used corresponds to native residues N171 to S391. C177, found in the native sequence and forming a disulfide bridge with C121 of the PLL domain, was mutated to a serine in all constructs (dark red). Engineered disulfides are indicated as bold and underlined red letters. The BiP signal peptide is cleaved between SLG // AT where // is the cleavage site.

#### BiP-FgB-linker-GAIN_WT-linker-10His-Spytag (AFM)

MKLCILLAVVAFVGLSLGATNEEGFFFSARGHRPLDGSGSGSGSAGTGSGNASVDMSELKRDLQLLSQFLKHPQKASRRPSAAPASQQLQSLESKLTSVRFMGDMVSFEEDRINATVWKLQPTAGLQDLHIHSRQEEEQSEIMEYSVLLPRTLFQRTKGRSGEAEKRLLLVDFSSQALFQDKNSSQVLGEKVLGIVVQNTKVANLTEPVVLTFQHQLQPKNVTLQCVFWVEDPTLSSPGHWSSAGCETVRRETQTSCFCNHLTYFAVLMVSGSGSGSGSAGTGSGHHHHHHHHHHGSAHIVMVDAYKPTK

#### BiP-FgB-linker-GAIN_<u>SS_h2l10</u>-linker-10His-Spytag (AFM)

MKLCILLAVVAFVGLSLGATNEEGFFFSARGHRPLDGSGSGSGSAGTGSGNASVDMSELKRDLQLLSQFLKHPQKASRRPSAAPA**C**QQLQSLESKLTSVRFMGDMVSFEEDRINATVWKLQPTAGLQDLHIHSRQEEEQSEIMEYSVLLPRTLFQRTKGRSGEAEKRLLLVDFSSQALFQDKNSSQVLGEKVLGIVVQNTKVANLTEPVVLTFQHQLQPKNVTLQCVFWV**C**DPTLSSPGHWSSAGCETVRRETQTSCFCNHLTYFAVLMVSGSGSGSGSAGTGSGHHHHHHHHHHGSAHIVMVDAYKPTK

#### BiP-FgB-linker-GAIN_<u>SS_h1h2</u>-linker-10His-Spytag (AFM)

MKLCILLAVVAFVGLSLGATNEEGFFFSARGHRPLDGSGSGSGSAGTGSGNAS**C**DMSELKRDLQLLSQFLKHPQKASRRPSAAPASQQLQSLESKLT**C**VRFMGDMVSFEEDRINATVWKLQPTAGLQDLHIHSRQEEEQSEIMEYSVLLPRTLFQRTKGRSGEAEKRLLLVDFSSQALFQDKNSSQVLGEKVLGIVVQNTKVANLTEPVVLTFQHQLQPKNVTLQCVFWVEDPTLSSPGHWSSAGCETVRRETQTSCFCNHLTYFAVLMVSGSGSGSGSAGTGSGHHHHHHHHHHGSAHIVMVDAYKPTK

#### BiP-FgB-linker-GAIN_<u>SS_h2l10-h1h2</u>-linker-10His-Spytag (AFM)

MKLCILLAVVAFVGLSLGATNEEGFFFSARGHRPLDGSGSGSGSAGTGSGNAS**C**DMSELKRDLQLLSQFLKHPQKASRRPSAAPA**C**QQLQSLESKLT**C**VRFMGDMVSFEEDRINATVWKLQPTAGLQDLHIHSRQEEEQSEIMEYSVLLPRTLFQRTKGRSGEAEKRLLLVDFSSQALFQDKNSSQVLGEKVLGIVVQNTKVANLTEPVVLTFQHQLQPKNVTLQCVFWV**C**DPTLSSPGHWSSAGCETVRRETQTSCFCNHLTYFAVLMVSGSGSGSGSAGTGSGHHHHHHHHHHGSAHIVMVDAYKPTK

#### BiP-GAIN_<u>SS_h2l10-h1h2</u>-linker-10His (X-ray crystallography)

MKLCILLAVVAFVGLSLGATNASCDMSELKRDLQLLSQFLKHPQKASRRPSAAPACQQLQSLESKLTCVRFMGDMVSFEEDRINATVWKLQPTAGLQDLHIHSRQEEEQSEIMEYSVLLPRTLFQRTKGRSGEAEKRLLLVDFSSQALFQDKNSSQVLGEKVLGIVVQNTKVANLTEPVVLTFQHQLQPKNVTLQCVFWVCDPTLSSPGHWSSAGCETVRRETQTSCFCNHLTYFAVLMVSGSGSHHHHHHHHHH

#### ybbR-6His-ELP-FLN-SpyCatcher (AFM surface handle)

MGTDSLEFIASKLAHHHHHHWGSGHGVGVPGMGVPGVGVPGVGVPGVGVPGVGVPGVGVPGVGVPGVGVPGEGVPGEGVPGVGVPGMGVPGVGVPGVGVPGVGVPGVGVPGVGVPGVGVPGVGVPGEGVPGEGVPGVGVPGMGVPGVGVPGVGVPGVGVPGVGVPGVGVPGVGVPGVGVPGEGVPGEGVPGWPSGSADPEKSYAEGPGLDGGECFQPSKFKIHAVDPDGVHRTDGGDGFVVTIEGPAPVDPVMVDNGDGTYDVEFEPKEAGDYVINLTLDGDNVNGFPKTVTVKPAPGSGSGSGSVDTLSGLSSEQGQSGDMTIEEDSATHIKFSKRDEDGKELAGATMELRDSSGKTISTWISDGQVKDFYLYPGKYTFVETAAPDGYEVATAITFTVNEQGQVTVNGKATKGDAHI

#### SdrG-FLN-ELP-6His-ybbr (AFM cantilever handle)

MGTEQGSNVNHLIKVTDQSITEGYDDSDGIIKAHDAENLIYDVTFEVDDKVKSGDTMTVNIDKNTVPSDLTDSFAIPKIKDNSGEIIATGTYDNTNKQITYTFTDYVDKYENIKAHLKLTSYIDKSKVPNNNTKLDVEYKTALSSVNKTITVEYQKPNENRTANLQSMFTNIDTKNHTVEQTIYINPLRYSAKETNVNISGNGDEGSTIIDDSTIIKVYKVGDNQNLPDSNRIYDYSEYEDVTNDDYAQLGNNNDVNINFGNIDSPYIIKVISKYDPNKDDYTTIQQTVTMQTTINEYTGEFRTASYDNTIAFSTSSGQGQGDLPPEGSGSGSGSADPEKSYAEGPGLDGGECFQPSKFKIHAVDPDGVHRTDGGDGFVVTIEGPAPVDPVMVDNGDGTYDVEFEPKEAGDYVINLTLDGDNVNGFPKTVTVKPAPGSGSGSHGVGVPGMGVPGVGVPGVGVPGVGVPGVGVPGVGVPGVGVPGVGVPGEGVPGEGVPGVGVPGMGVPGVGVPGVGVPGVGVPGVGVPGVGVPGVGVPGVGVPGEGVPGEGVPGVGVPGMGVPGVGVPGVGVPGVGVPGVGVPGVGVPGVGVPGVGVPGEGVPGEGVPGWRGHHHHHHGSDSLEFIASKLA

## References

1. M. Lerche et al., iScience. 23, 100907 (2020).

2. S. Seetharaman, S. Etienne-Manneville, Biology of the Cell. 110, 49–64 (2018).

3. V. Vogel, M. Sheetz, Nat Rev Mol Cell Biol. 7, 265–275 (2006).

4. Y. Jiang, X. Yang, J. Jiang, B. Xiao, Trends in Biochemical Sciences. 46, 472–488 (2021).

5. R. Gnanasambandam, P. A. Gottlieb, F. Sachs, in Current Topics in Membranes, P. A. Gottlieb, Ed. (Academic Press, 2017; https://www.sciencedirect.com/science/article/pii/S1063582316300461), vol. 79 of *Piezo Channels*, pp. 275–307.

6. F. Bassilana, M. Nash, M.-G. Ludwig, Nat Rev Drug Discov. 18, 869–884 (2019).

7. A. Vizurraga, R. Adhikari, J. Yeung, M. Yu, G. G. Tall, J Biol Chem. 295, 14065–14083 (2020).

8. D. Araç, K. Leon, in GPCRs, B. Jastrzebska, P. S.-H. Park, Eds. (Academic Press, 2020; https://www.sciencedirect.com/science/article/pii/B9780128162286000027), pp. 23–41.

9. I. Liebscher et al., The FEBS Journal. **n/a,** doi:10.1111/febs.16258.

10. J. Hamann et al., Pharmacol Rev. 67, 338–367 (2015).

11. C. Wilde, J. Mitgau, T. Suchý, T. Schöneberg, I. Liebscher, Am J Physiol Cell Physiol. 322, C1047–C1060 (2022).

12. J. Mitgau et al., Front Cell Dev Biol. 10, 873278 (2022).

13. J. Yeung et al., Proc Natl Acad Sci U S A. 117, 28275–28286 (2020).

14. D. Araç et al., EMBO J. 31, 1364–1378 (2012).

15. N. Scholz et al., Nature. 615, 945–953 (2023).

16. B. Zhu et al., J. Biol. Chem. 294, 19246–19254 (2019).

17. I. Liebscher, T. Schöneberg, D. Thor, Signal Transduct Target Ther. 7, 227 (2022).

18. X. Barros-Álvarez et al., Nature. 604, 757–762 (2022).

19. P. Xiao et al., Nature. 604, 771–778 (2022).

20. Y.-Q. Ping et al., Nature. 604, 763–770 (2022).

21. X. Qu et al., Nature. 604, 779–785 (2022).

22. C. Mao et al., Molecular Cell. 84, 570–583.e7 (2024).

23. G. Beliu et al., Mol Cell. 81, 905–921.e5 (2021).

24. G. S. Salzman et al., Proc Natl Acad Sci U S A. 114, 10095–10100 (2017).

25. S. P. Kordon et al., Nat Commun. 15, 10545 (2024).

26. L. F. Milles, K. Schulten, H. E. Gaub, R. C. Bernardi, Science. 359, 1527–1533 (2018).

27. V. Vogel, Annu Rev Physiol. 80, 353–387 (2018).

28. C. Schoeler et al., Nano Lett. 15, 7370–7376 (2015).

29. B. Kuhlman, P. Bradley, Nat Rev Mol Cell Biol. 20, 681–697 (2019).

30. M. Marfoglia, L. Guirardel, M. Pedraza, A. Chatzi Souleiman, P. Barth, AlloPool: Predicting Protein Allostery using Deep Learning (2024), p. 2024.11.01.621466, , doi:10.1101/2024.11.01.621466. (see companion paper)

31. G. S. Salzman et al., Neuron. 91, 1292–1304 (2016).

32. C. Fu et al., Nano Lett. 23, 9179–9186 (2023).

33. Piconewton Forces Mediate GAIN Domain Dissociation of the Latrophilin-3 Adhesion GPCR | Nano Letters, (available at https://pubs.acs.org/doi/full/10.1021/acs.nanolett.3c03171).

34. M. Rief, M. Gautel, F. Oesterhelt, J. M. Fernandez, H. E. Gaub, Science. 276, 1109–1112 (1997).

35. S. P. Ng et al., J Mol Biol. 350, 776–789 (2005).

36. A. F. Oberhauser, C. Badilla-Fernandez, M. Carrion-Vazquez, J. M. Fernandez, J Mol Biol. 319, 433–447 (2002).

37. Z. Liu et al., Nano Lett. 22, 179–187 (2022).

38. D. J. Brockwell et al., Nat Struct Biol. 10, 731–737 (2003).

39. M. Carrion-Vazquez et al., Nat Struct Biol. 10, 738–743 (2003).

40. Y. D. Li, G. Lamour, J. Gsponer, P. Zheng, H. Li, Biophys J. 103, 2361–2368 (2012).

41. H. Dietz, F. Berkemeier, M. Bertz, M. Rief, Proc Natl Acad Sci U S A. 103, 12724–12728 (2006).

42. Direct Comparison of Lysine versus Site-Specific Protein Surface Immobilization in Single-Molecule Mechanical Assays** - Liu - 2023 - Angewandte Chemie International Edition - Wiley Online Library, (available at https://onlinelibrary.wiley.com/doi/10.1002/anie.202304136).

43. F. Li, S. D. Redick, H. P. Erickson, V. T. Moy, Biophys J. 84, 1252–1262 (2003).

44. F. Rico, L. Gonzalez, I. Casuso, M. Puig-Vidal, S. Scheuring, Science. 342, 741–743 (2013).

45. W. R. Gordon et al., Dev Cell. 33, 729–736 (2015).

46. V. C. Luca et al., Science. 355, 1320–1324 (2017).

47. B. Shergill, L. Meloty-Kapella, A. A. Musse, G. Weinmaster, E. Botvinick, Developmental Cell. 22, 1313–1320 (2012).

48. J. Zhu, J. Wang, W. Han, D. Xu, Nat Commun. 13, 1661 (2022).

49. H. M. Stoveken, A. G. Hajduczok, L. Xu, G. G. Tall, Proc Natl Acad Sci U S A. 112, 6194–6199 (2015).

50. R. Luo et al., Proc Natl Acad Sci U S A. 108, 12925–12930 (2011).

51. J. Jumper et al., Nature. 596, 583–589 (2021).

52. Influence of Fluorination on Single-Molecule Unfolding and Rupture Pathways of a Mechanostable Protein Adhesion Complex | Nano Letters, (available at https://pubs.acs.org/doi/10.1021/acs.nanolett.0c04178).

53. R. Merkel, P. Nassoy, A. Leung, K. Ritchie, E. Evans, Nature. 397, 50–53 (1999).

54. E. Evans, K. Ritchie, Biophysical Journal. 72, 1541–1555 (1997).

55. O. K. Dudko, Q Rev Biophys. 49, e3 (2016).

56. D. Ok, H. G, S. A, Proceedings of the National Academy of Sciences of the United States of America. 105 (2008), doi:10.1073/pnas.0806085105.

57. A. D’Arcy, F. Villard, M. Marsh, Acta Cryst D. 63, 550–554 (2007).

58. W. Kabsch, J Appl Cryst. 26, 795–800 (1993).

59. A. J. McCoy et al., J Appl Cryst. 40, 658–674 (2007).

60. P. Emsley, K. Cowtan, Acta Cryst D. 60, 2126–2132 (2004).

61. P. D. Adams et al., Acta Cryst D. 66, 213–221 (2010).

62. V. B. Chen et al., Acta Cryst D. 66, 12–21 (2010).

63. J. Dauparas, et al., Science. 378, 49–56 (2022).

64. N. Ollikainen, R. M. de Jong, T. Kortemme, PLoS Comput Biol. 11, e1004335 (2015).

65. M. J. Abraham et al., SoftwareX. 1–2, 19–25 (2015).

66. Update of the CHARMM All-Atom Additive Force Field for Lipids: Validation on Six Lipid Types | The Journal of Physical Chemistry B, (available at https://pubs.acs.org/doi/10.1021/jp101759q).

67. P. Mark, L. Nilsson, J. Phys. Chem. A. 105, 9954–9960 (2001).

68. U. Essmann et al., The Journal of Chemical Physics. 103, 8577–8593 (1995).

69. E.W. Dijkstra Archive: A Short Introduction to the Art of Programming (EWD 316), (available at https://www.cs.utexas.edu/users/EWD/transcriptions/EWD03xx/EWD316.html).

70. P. Wernet et al., Science. 304, 995–999 (2004).

71. S. Jo, T. Kim, V. G. Iyer, W. Im, J Comput Chem. 29, 1859–1865 (2008).

72. J. Lee et al., J Chem Theory Comput. 12, 405–413 (2016).

73. J. Huang et al., Nat. Methods. 14, 71–73 (2017).

